# Open-ST: High-resolution spatial transcriptomics in 3D

**DOI:** 10.1101/2023.12.22.572554

**Authors:** Marie Schott, Daniel León-Periñán, Elena Splendiani, Leon Strenger, Jan Robin Licha, Tancredi Massimo Pentimalli, Simon Schallenberg, Jonathan Alles, Sarah Samut Tagliaferro, Anastasiya Boltengagen, Sebastian Ehrig, Stefano Abbiati, Steffen Dommerich, Massimiliano Pagani, Elisabetta Ferretti, Giuseppe Macino, Nikos Karaiskos, Nikolaus Rajewsky

## Abstract

Spatial transcriptomics (ST) methods have been developed to unlock molecular mechanisms underlying tissue development, homeostasis, or disease. However, there is a need for easy-to-use, high-resolution, cost-efficient, and 3D-scalable methods. Here, we report Open-ST, a sequencing-based, open-source experimental and computational resource to address these challenges and to study the molecular organization of tissues in 3D. In mouse brain, Open-ST captured transcripts at subcellular resolution and reconstructed cell types. In primary tumor and patient-matched healthy/metastatic lymph nodes, Open-ST captured the diversity of immune, stromal and tumor populations in space. Distinct cell states were organized around cell-cell communication hotspots in the tumor, but not the metastasis. Strikingly, the 3D reconstruction and multimodal analysis of the metastatic lymph node revealed spatially contiguous structures not visible in 2D and potential biomarkers precisely at the 3D tumor/lymph node boundary. We anticipate Open-ST to accelerate the identification of spatial molecular mechanisms in 2D and 3D.

## Introduction

Recent years have witnessed a massive increase in the development and application of spatially resolved transcriptomics (ST) methods ^1^. Unlike standard single-cell methods, ST retains the spatial context of the captured transcriptome and thus allows the direct observation of the arrangements of cells and their interactions in tissue space. These data may be of fundamental importance for understanding molecular mechanisms in health and also critical for identifying and targeting the molecular origins of diseases ^2–4^. For example, tumor microenvironment interactions or the spatial structure of lymph nodes are critical to understand function ^5,6^. Moreover, ST may avoid biases introduced by single-cell dissociation, which depletes certain cell types and activates stress pathways ^7–9^.

Commercially available ST technologies that provide non-targeted capture of transcriptomes are limited by their relatively high costs and/or limited resolution; these include Visium (10X Genomics), CurioSeeker (Curio Bioscience), and Stereo-seq (BGI)^10^. Probe-based (targeted) methods, including CosMx Spatial Molecular Imager, GeoMx Digital Spatial Profiler (Nanostring), Molecular Cartography (Resolve Biosciences), Xenium In Situ (10X Genomics) target a predesigned panel of genes and are therefore not suited for unbiased discovery or spatial genotyping ^11,12,13^. Other spatial technologies developed in non-commercial settings can be challenging to implement (XYZ-seq, Pixel-seq, Seq-Scope, Sci-Space, High-Definition Spatial Transcriptomics, Slide-seqV2, DBiT-seq) ^14,15,16,17,18,19,20^. Finally, although cells operate and communicate in 3D, building up functional tissues and organs, no end-to-end platform currently exists to generate and computationally analyze ST in 3D.

Here, we present Open-ST, a spatial transcriptomics method for fresh-frozen tissue samples that enables efficient capture of polyadenylated transcripts at subcellular resolution and includes open-source software for seamless data processing and analysis in 3D. Open-ST operates by converting Illumina flow cells into capture areas, an approach previously implemented in Seq-Scope ^16^. Our method encompasses several key enhancements.

First, we use patterned flow cell technology to create densely barcoded areas that capture RNA from a tissue section at a capture spot resolution of ∼0.6 μm. To control the fragmentation of the flow cell into distinct capture areas, we provide a 3D printable cutting guide. Our simplified library preparation only requires standard lab equipment, and comes with a cost of <150 € per total 12 mm^2^ capture area. The total costs of a standard sample (3 x 4 mm, 400M sequencing reads, ∼100,000 cells, ∼1000 UMIs/cell) are a few hundred Euros, primarily driven by the sequencing costs.

Compared to other sequencing-based technologies, Open-ST required the least sequencing depth to obtain an equivalent amount of transcriptomic information, making it cost-effective. It only requires access to a sequencing facility and otherwise standard lab equipment. Open-ST is scalable, as a single researcher can prepare 10-15 libraries in three days starting from prepared capture areas, and versatile, as capture area size can be adjusted within the limitations of the flow cell size (∼80x7mm for NovaSeq6000 S4). Our Hematoxylin and Eosin (H&E) imaging pipeline, which we optimized for frozen samples, produces high-resolution images from the same section, which Open-ST uses for cell segmentation and integrates with transcriptomic data. Open-ST generated 2D data are robust enough to be computationally integrated into 3D (“virtual tissue blocks”). The single-cell segmentation, subcellular resolution, and 3D tissue reconstruction and interrogation capabilities are powered by a stack of computational tools conceived as a modular software package specifically designed for Open-ST data. We built our tools within a human-in-the-loop paradigm, enabling data interactivity with GUI tools and efficient data structures. Our software to interrogate is agnostic to the orientation of the original slices and can be used to discover 3D molecular patterns and potential biomarkers. The entire software stack is open-source.

We demonstrate that Open-ST recapitulates cell types and marker genes of several tissues (mouse and human) with subcellular precision. Embryonic mouse head and adult mouse hippocampus were used to carefully benchmark the precision, sensitivity, and spatial resolution of RNA capture due to the high availability of published gene expression data (RNA-seq, *in-situ* hybridization, ST, etc.) and the possibility to maintain RNA quality by controlling sample handling and timing.

To demonstrate the capability of Open-ST to profile clinically relevant tissues encompassing drastically different morphologies and cell sizes (from small immune cells to ∼100x larger adipocytes), we profiled patient-matched samples from human head and neck squamous cell carcinoma (HNSCC). This cancer type has been shown to have a high diversity of transcriptional profiles and cell type composition, whose relative abundance, spatial organization and interaction have implications in survival and therapy response ^21–24^. Across all 21 human sections processed with Open-ST, we reproducibly captured, at moderate sequencing depths, a median of ∼600-2,000 spatially mapped transcripts per cell, covering more than 25,000 genes. We also show that our libraries were sequenced far from saturation, and that resequencing increased the molecular counts.

For the human metastatic lymph node, we applied Open-ST to obtain H&E staining images and ST from serial sections spanning 350 μm. We constructed a 3D virtual tissue block with more than a million sequenced cells and 851 million transcripts embedded in the H&E stainings. In the patient-matched primary tumor, Open-ST recapitulated the transcriptomic identity of stromal, immune and tumor cell types from the metastatic tissue. In particular, sub-clustering identified 10 tumor cell subtypes present in both the primary and the metastatic tumor. Strikingly, these subtypes were spatially patterned in the primary tumor, but not in the metastatic tissue. They included proliferative, inflammatory, keratinizing and invasive phenotypes and correlated strongly to the spatial localization of cell-cell communication hotspots, computationally predicted by ligand-receptor analyses. The 3D virtual tissue block of the metastatic tumor in the lymph node allowed us to identify potential biomarkers in 3D. Specifically, we detected a spatially organized cholesterol biosynthesis signature and a population of macrophages at the 3D boundary between tumor and lymphoid tissue.

Due to its ease-of-use, cost-effectiveness, and wide applicability, we envision Open-ST to become a valuable method for diverse spatial -omics studies. To aid researchers in implementing and using Open-ST, we have set up an online resource with detailed descriptions of all experimental and computational protocols/software (https://rajewsky-lab.github.io/openst, Figure 1A).

**Figure 1:**
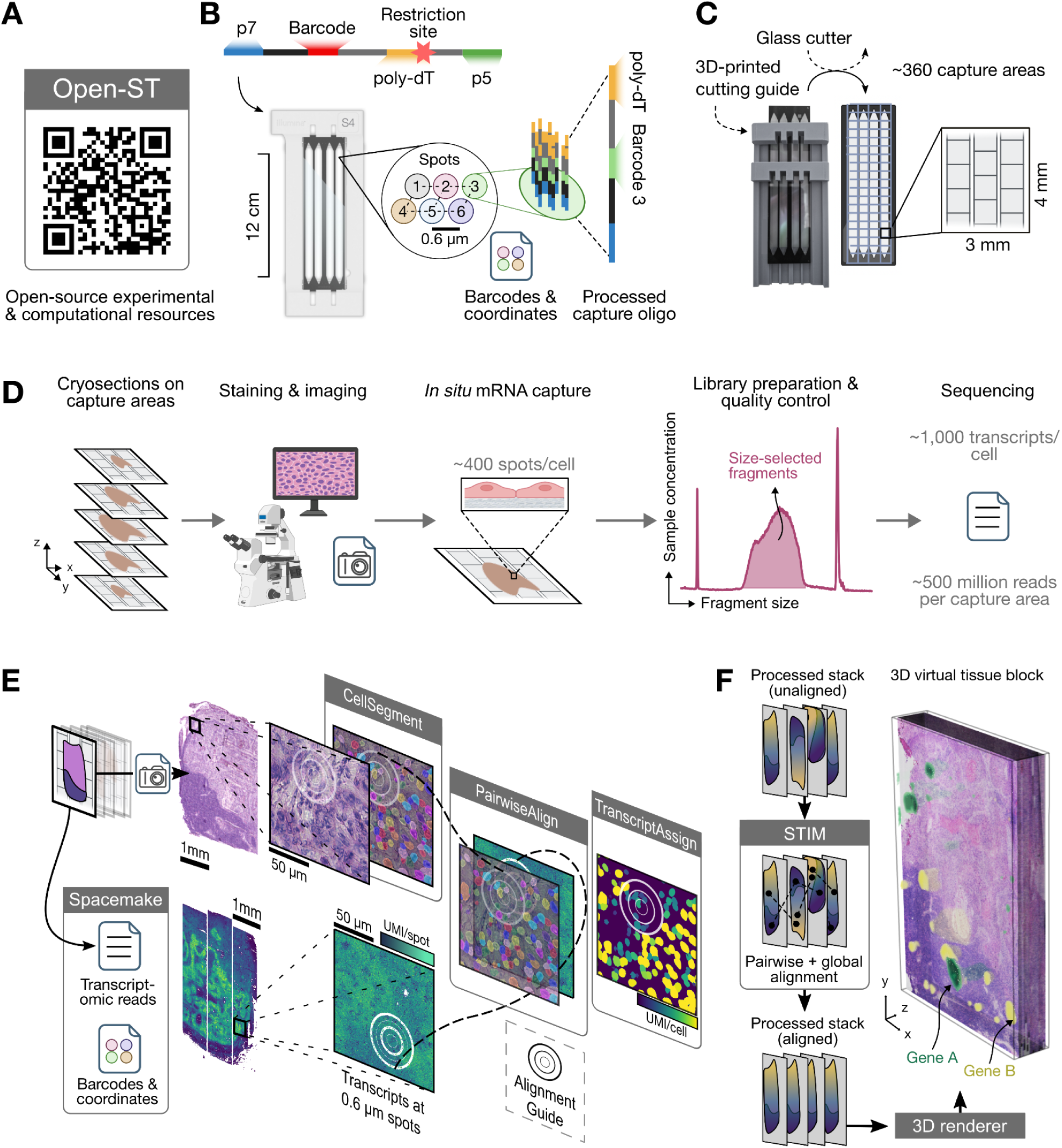
Open-ST workflow for high-resolution spatial transcriptomics of segmented single-cells in 2D or 3D. **(A)** All experimental and computational resources are open-source and available under https://rajewsky-lab.github.io/openst (QR code). **(B)** Sequencing designed oligos in patterned Illumina flow cells (here, NovaSeq6000 S4) allows barcode registration in regularly spaced spots (left). High-density spots with clonal oligonucleotides are processed to allow capture of poly-adenylated RNA (right). Flow cell image courtesy of Illumina, Inc. **(C)** The opened and processed flow cell is placed on our custom 3D-printable device that allows guided cutting to produce ∼360 capture areas of 3x4 mm^2^ at ∼45€ each (Methods). The capture area is composed of ∼1 mm^2^ tiles across 3 columns. p5/p7: Illumina adapters. (**D**) Transcriptomic and H&E imaging data are generated from the same fresh-frozen tissue section. Optimized RNA capture conditions and a single-amplification library preparation result in high library complexity (Methods). FU, fluorescent units. (**E**) Tissue morphological information (imaging) is integrated with spatial transcriptomics data from single sections with our open-source `openst` Python package, including code for automatic cell segmentation, pairwise alignment of modalities, and quantification of transcripts (unique molecular identifiers, UMI) to segmented cells (Methods). **(F)** Serial sections can be used for three-dimensional reconstruction of tissue histology and transcriptome, using STIM (left). Imaging and transcriptomics data can be visualized and interrogated as a 3D virtual tissue block using any 3D rendering engine (right) (Methods). The smoothed, volumetric rendering of two genes (Gene A: *S100A7*; Gene B: *FDCSP*) is shown for illustration purposes.

## Results

### 1. The Open-ST workflow

To generate the mRNA capture areas, we leverage Illumina’s sequencing-by-synthesis technology, using a custom sequencing recipe (Supplemental information 1). In brief, we register spatial barcode sequences and their associated (x, y) coordinates on the flow cell, by sequencing oligos which comprise unique 32-nucleotide barcodes, appropriate adapters, and a poly-dT region (Figure 1B, Methods). Bridge amplification at the start of sequencing generates densely packed spots, each containing thousands of clonal oligonucleotides with a unique barcode sequence. To maximize information retrieval and resolution, Open-ST employs the patterned NovaSeq6000 flow cells for spot generation, which use regularly spaced nanowells with a center-to-center distance of ∼0.6 μm. Patterning reduces the probability of acquiring mixed-barcode signals within a single spot and yields higher spot density compared to non-patterned flow cells ^25^.

Following the sequencing of the designed capture oligo barcodes, we process the oligonucleotides to allow polyA transcript capture and open the flow cell (Methods). Our custom, 3D-printable tool facilitates the cutting of the opened flow cell into small capture areas, whilst preventing surface scratches (Figure 1C, Methods). The dimensions of the capture area can be chosen based on the experimental design, with a maximum size of 7 x 80 mm as limited by the sequenced area of the flow cell (Figure S1A). At 3 x 4 mm, around 360 capture areas can be made from one NovaSeq S4 flow cell, each at around 45€. These showed high spatial regularity of capture spots with only a few artifacts (Figure S1B, C), and barcode sequences with the expected structure (Figure S1D).

Open-ST allows the analysis of tissue morphology with a high-quality H&E staining and localized capture of RNA transcripts from the same cryo-section (Figure 1D). Pepsin and hybridization buffer (2x saline-sodium citrate, SSC) were combined in one solution to promote the simultaneous tissue permeabilization and RNA capture, by reducing electrostatic repulsion of the single-stranded DNA and RNA molecules. A qPCR assay is used to determine optimal permeabilization conditions for maximum mRNA capture. Implementation of a qPCR-based quality control during library preparation allows us to determine the optimal number of PCR cycles to avoid over- or under-amplification of the library. Additionally, the one-step library amplification prevents PCR amplification bias or sample losses due to bead purification between PCR reactions (Figure S1E, F, Methods).

The availability of morphological features and molecular readouts from the same tissue section enables efficient cell segmentation (Figure 1E). The H&E raw images are automatically preprocessed (Figure S1G) and then segmented into single cells using a fine-tuned Cellpose model (Figure S1H, Methods) ^26^. Following nuclei segmentation, radial extension of nuclei boundaries adds cytoplasmic context to yield approximate cell boundaries (Figure S1G,H, Methods). Using the barcode coordinates from the first sequencing run, the transcriptomic reads are processed and mapped in tissue space using Spacemake ^27^. Subsequently, the flow cell circular marks, visible in both the imaging and spatial transcriptome modalities, are used as alignment guides. These guides are automatically detected for unsupervised pairwise alignment, resulting in registration of the imaging and sequencing data with an accuracy of ∼1 μm per tile (Figure S1I, Methods). Our segmentation and alignment protocol unbiasedly adapts to tissues with heterogeneous cell sizes and densities, and automatically excludes tissue areas without cells from downstream analyses (Figure S1H, Methods). Image preprocessing and fine-tuning of the segmentation model increased the precision of the segmentation, as evidenced by benchmarking against a manual segmentation (Figure S1J). Consequently, Open-ST data can be analyzed at the single-cell level and integrated with the (histological) staining data.

Given the efficient transcript capture and our integrated computational pipeline, Open-ST is suitable for 3D spatial reconstruction of any tissue. Here, we employ it for the reconstruction of a metastatic lymph node, integrating the H&E staining and gene expression data (Figure 1F). The serial sections were aligned using the open-access Spatial Transcriptomics Imaging Framework (STIM), resulting in a 3D spatial representation of the tissue ^28^. The imaging and transcriptomics data can be visualized and interrogated as a 3D virtual tissue block using any 3D rendering software. In particular, we used an open-source graphical user interface tool optimized for scientific visualization ^29^.

### 2. Open-ST robustly captures transcripts with high efficiency

We successfully applied our method in diverse tissues: embryonic mouse head, adult mouse hippocampus, human primary tumor (HNSCC), and the patient-matched healthy and metastatic lymph nodes.

All samples exhibited a high percentage of transcripts mapping uniquely to the genome (65-78%, Table S1), with only a small percentage of reads consisting of ribosomal RNA (2.5-15.3%, Table S1). A mean of ∼55% of the reads that map to genic regions were assigned to a spatial barcode from the first sequencing run (Table S1). Open-ST retains more reads when benchmarked against alternative solutions, and demonstrates consistency across samples (Figure S2A). We found that the percentage of barcode reads from the first sequencing detected in the transcriptome library was related to tissue coverage and cell density (Figure S2B). Outliers may represent segmentation errors, capture of ambient RNA in regions without cells, or areas with cells but no spatial barcodes due to capture area irregularities. Figure S2C illustrates the breakdown of the total read numbers for a representative sample after alignment, spatial mapping and deduplication.

In a sagittal E13 mouse brain section, sequenced at a depth of 478 million reads, we segmented 49,048 cells, capturing a total of 21,609 genes (Figure 2A). Open-ST’s efficient RNA capture resulted in a median of 621 genes and 880 UMIs (Figure 2B, Table S2), with 42% of the cells containing over 1,000 transcripts. We explored the contribution of all detected genes to transcriptomic information, finding that across the entire dataset, 10,000 genes account for 95% of the captured transcriptomic information. Similarly, the median contributions of background transcripts were 15.4% for ribosomal proteins and 3.5% for mitochondria-encoded transcripts (Figure S2D). At the same time, Open-ST resulted in rich quantification of all genes, ranging from the most abundant transcript, *mt-Cytb*, which accounted for ∼1% of all UMIs and was present in almost all cells, to lowly expressed but highly specific gene markers (Figure S2E). Thus, a notable portion of detected genes is well-represented across all cells.

**Figure 2:**
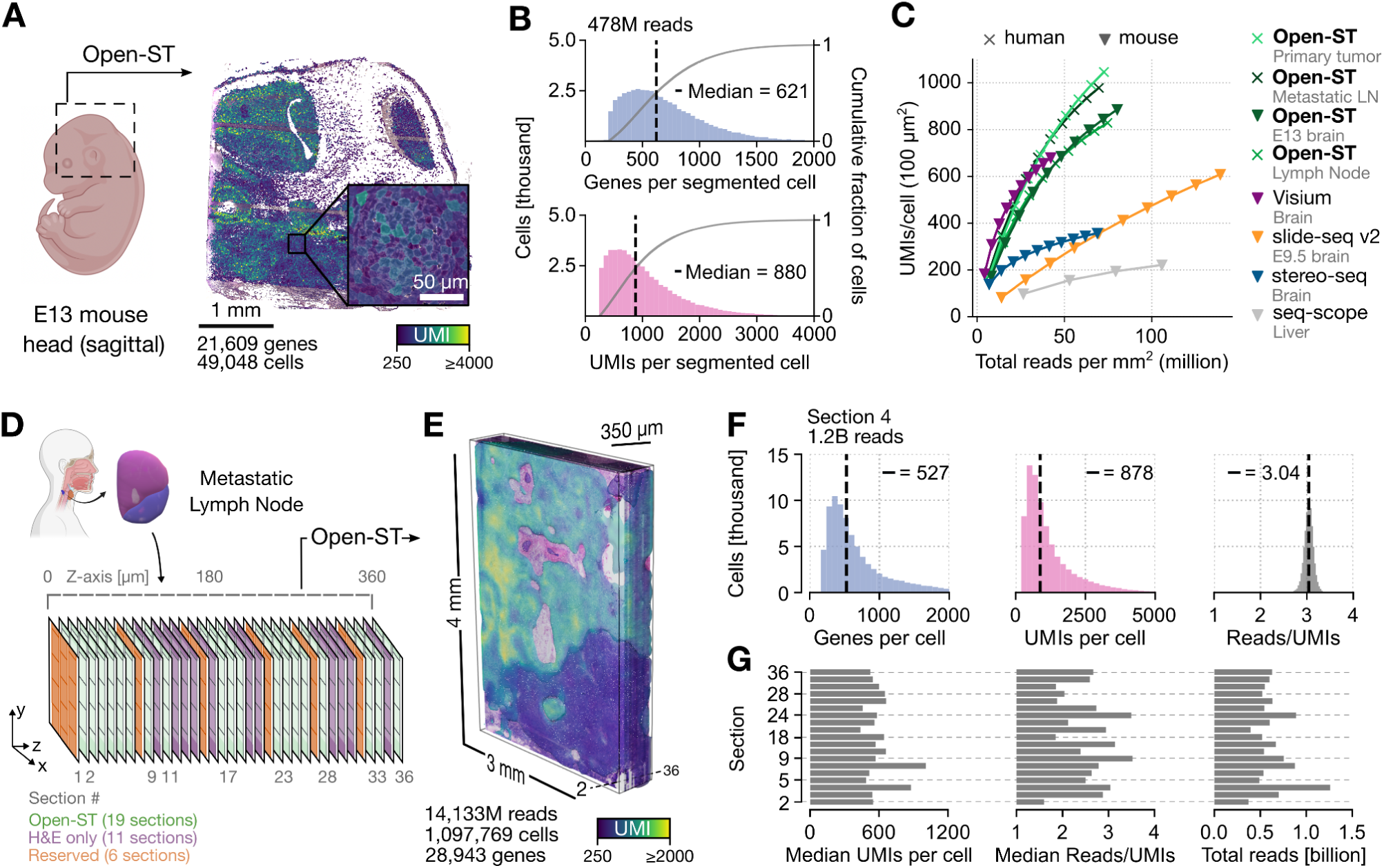
Open-ST robustly captures transcripts with high efficiency. **(A)** Spatial distribution of UMI counts per segmented cell in an Open-ST processed E13 mouse head sagittal section. **(B)** Distribution of genes and UMIs per segmented cell; sample as in (A). (**C)** Open-ST produces the highest transcript capture per pseudo-cell (UMIs/100µm^2^) for the same number of sequencing reads compared to alternative sequencing-based spatial transcriptomic methods (Methods). **(D)** Experimental design for processing a human metastatic lymph node (LN) with Open-ST. 10 μm thick sections were processed and profiled with Open-ST, only H&E stained, or reserved for validations. **(E)** Aligned transcriptomic and imaging data; rendering of transcript capture colored by UMI counts is overlaid on a 3D virtual tissue block (Methods). **(F)** Distributions of UMIs and gene counts per segmented cell for section 4, which was sequenced the deepest due to its low reads/UMIs ratio. Median values are indicated. **(G)** Total reads, median UMIs and median reads/UMIs per segmented cell across all 19 sections show consistently high transcript capture and low ratio of sequencing reads to UMIs captured.

We assessed the library complexity of Open-ST data in comparison to existing sequencing-based ST methods by calculating UMI counts per pseudo-cell (100 µm^2^) relative to sequencing depth (Figure 2C, Methods). Open-ST consistently outperformed alternative solutions in capture efficiency for the same sequencing depth with the exception of 10X Visium, which demonstrated comparable performance. Importantly, capture efficiency was similar between Open-ST processed samples, despite their diverse cellular composition and variable RNA quality (RIN 6.7-8.9, Table S1). To further evaluate library complexity across technologies, we benchmarked the ratio of reads over UMIs as a function of genic reads (Figure S2F, Methods). Open-ST consistently showed the lowest ratio and scaled best with increasing read depth, together with the Slide-seqV2 sample. A low reads-to-UMIs ratio results from high initial transcript capture and efficient library amplification, and is vital for extracting new information through deeper sequencing. Given that library sequencing constitutes the largest fraction of the total cost per sample, Open-ST emerges as an affordable and highly efficient solution for a whole transcriptome sequencing approach.

### 3. Open-ST generates 3D virtual tissue blocks

To demonstrate robustness and reproducibility of Open-ST across multiple sections in a clinically relevant complex tissue, we processed a human metastatic lymph node. In this proof-of-principle, large-scale experiment, we obtained 10 μm thick sections spanning a tissue depth of 370 μm. Our experimental design consisted of profiling sections with Open-ST (19), H&E-staining only (11), and reserving for validation (6) (Figure 2D). In total, we obtained gene expression profiles of over 1 million cells across 19 sections, with high median capture of genes (313-624) and UMIs (438-1,008) per segmented cell (Figure S2H, Table S2). The samples also demonstrated a similar percentage of reads aligning to rRNA (<15%) and mapping uniquely to the genome (>65%) (Figure S2I, Table S1).

By leveraging STIM and The Visualization Toolkit (VTK), we aligned the serial sections processed with Open-ST to reconstruct and visualize a 3D virtual tissue block, a computational object combining transcriptomics and histological modalities (Figure S2J) ^28,30^. A smooth isosurface rendering of the spatially mapped UMI counts indicates the reproducible capture efficiency with a visible transcript enrichment in the tumor compartment (Figure 2E). section 4 was sequenced at a high depth (1.2B reads), exhibiting low ratio of reads/UMIs (median=3.04) and resulting in a median of 527 (878) genes (UMIs) per segmented cell (Figure 2F). The reads/UMIs ratio was consistently low and comparable across sections, demonstrating the reproducibility and cost-effectiveness of Open-ST (Figure 2G, S2G).

Taken together, Open-ST robustly captures transcripts across serial sections and requires low sequencing investment for new transcript discovery. Thus, Open-ST is well-suited for high-throughput studies such as 3D-transcriptome reconstruction of tissues/organs, or the analysis of tissues from patient cohorts.

### 4. Open-ST locally captures marker genes with high accuracy

We corroborated localized transcript capture of Open-ST by investigating spatial cluster distribution and marker gene localization in the highly organized E13 mouse head. Unbiased clusters were annotated based on literature-informed marker genes, using a developmental mouse atlas as a reference (Table S3, and see Methods for what follows in this section) ^31^. Coarse clustering of segmented Open-ST data reflects the major anatomical regions in the embryonic mouse head (Figure 3A, Figure S3A). Spatial expression of selected marker genes per cluster co-localize with the respective clusters (Figure 3B, S3B).

**Figure 3.**
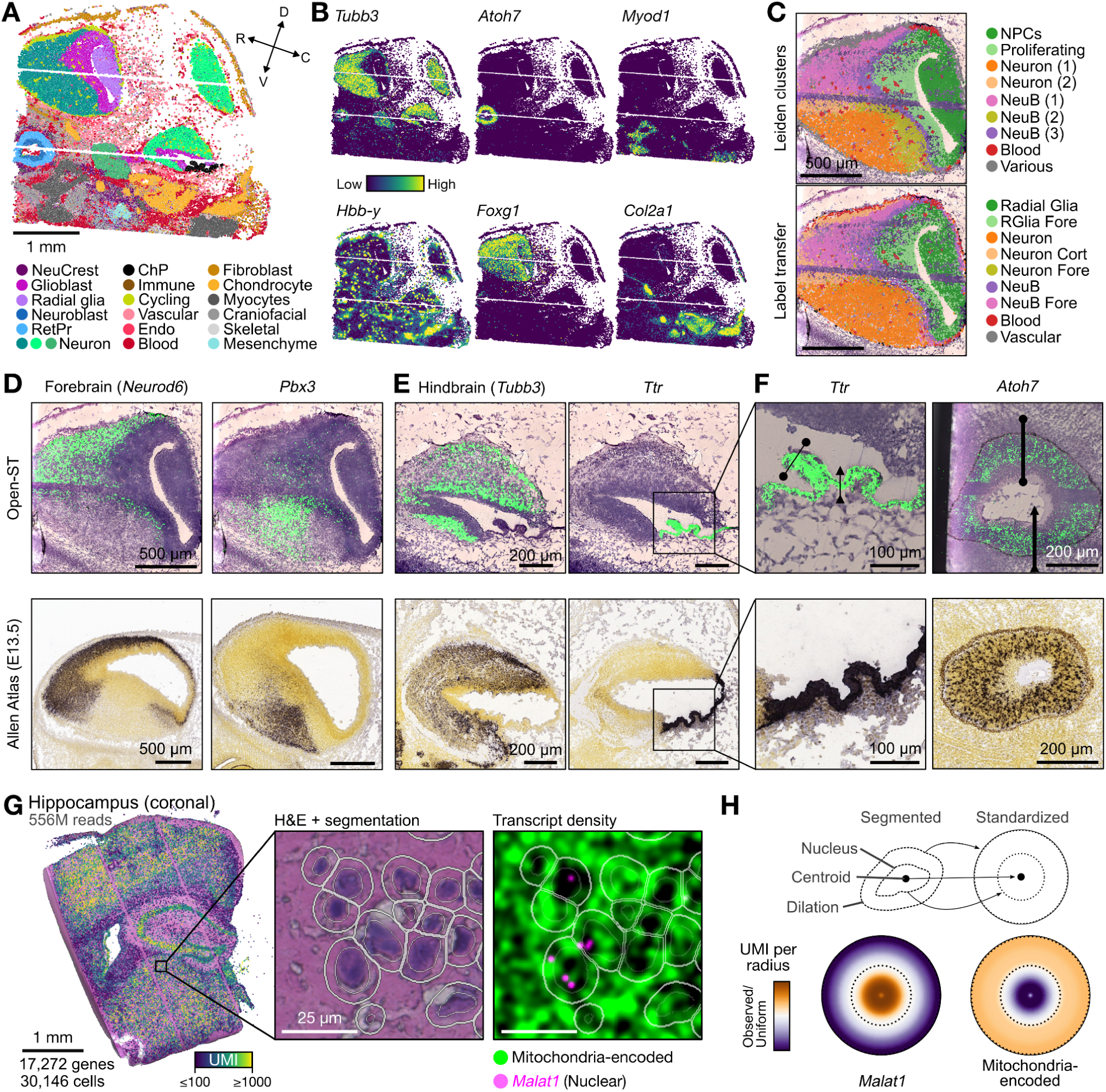
Open-ST captures transcripts at subcellular resolution. **(A)** Spatial distribution of cell types of an E13 mouse head sagittal section. D: dorsal, V: ventral, R: rostral, C: caudal. **(B)** Localization of selected marker genes colored by normalized expression. **(C)** Spatial distribution of E13 mouse forebrain cells colored by annotated Leiden cluster (top), and colored by the result of label transfer from an E13.5 single cell reference atlas (bottom) (Methods)^31^. **(D)** Localized gene capture of the *Neurod6* and *Pbx3* transcription factors in the E13 mouse forebrain profiled with Open-ST (top) compared to *in situ* hybridization images of the E13.5 mouse from the Allen Developing Mouse Brain Atlas (bottom) ^32,33^. High expression is colored in green (black) for Open-ST (Allen Atlas). **(E)** Localized gene capture in the E13 mouse hindbrain (*Tubb3*) and choroid plexus (*Ttr*) **(F)** Magnification on transcript density across two selected lines for *Ttr* in the choroid plexus; upper: transcript density in space is visualized as a 2D Kernel Density Estimate of the spot coordinates weighted by transcript counts; bottom: *in situ* hybridization images of E13.5 mouse (sagittal plane) from the Allen Developing Mouse Brain Atlas. **(G)** Subcellular transcript capture precision in an adult mouse coronal hippocampus section. Left: spatial distribution of UMIs per segmented cell reflects the tissue morphology; center: close-up of H&E segmentation with thin (thick) white lines indicating nuclear (cell) boundaries; right: nuclear-enriched *Malat1* and mitochondria-encoded transcript density is plotted onto the segmentation mask. **(H)** *Malat1* and mitochondria-encoded transcripts are enriched in the nucleus and cytoplasm, respectively. Observed over uniform UMI per radius (*n*_radii_ = 40), visualized on a standardized cell, quantified over 22,376 cells sampled from tissue regions where 15-30% area is covered by nuclei. Color mapping of expression is clipped between the 5th to 95th percentile of values (Methods).

We identified three neuronal clusters, each characterized by *Tbr1*, *Lhx9* or *Nefl* expression. A radial glia cluster, with cells mainly located in the dorsal forebrain, was identified by their expression of *Dmrta2*. The eye’s retina is set apart by localized expression of *Atoh7*, and the choroid plexus is characterized by *Ttr* expression in all cells of the cluster. Further examination led to the identification of fibroblasts (expressing collagens), chondrocytes (*Col11a1*, *Sox9*, *Hapln1*), myocytes (*Actc1, Mylpf*), mesenchymal (*Tsc22d1*) and endothelial cells (*Cd34* & *Kdr*). Coarse clustering revealed a group of cells with no identifiable markers. On these, we performed subclustering, and identified groups of cells that we termed neural crest (*Mdk*), craniofacial development (*Barx1*, *Sp7*), musculoskeletal development (*Kctd12*, *Igf12*), immune (*Selenop*, *Dab2*, *Ctsb*), and early vascular (*Akap12*, *Gja1*, *Cldn11*). The ‘blood’ cluster was identified based on its high expression of hemoglobin genes (e.g. *Hbb-y*). We found a cluster highly expressing the cell cycle gene *Ccnd2*, with transcriptomic distance closest to the neuron clusters, that we termed ‘cycling’. This cluster was in spatial proximity to the ‘blood’ cluster, which could explain the contaminating expression of hemoglobin genes (Figure S3B).

The capability of Open-ST to capture rich transcriptomic information allowed us to achieve more refined clustering of the segmented cells of the fore-, mid-, and hindbrain (Figure 3C, S4B, D, E, G, H). Cell type diversity and spatial distribution of cells in the forebrain resembled that obtained by integrating and transferring cell type labels from a single-cell reference atlas (see Methods) (Figure 3C). This congruence extends to marker genes identified from each unbiased cluster in our Open-ST data, demonstrating consistency with the respective populations in the reference (Figure S4B). This substantiates how the clustering of segmented Open-ST data provides labels akin to those obtained through the clustering of single-cell RNA sequencing data.

To investigate the localization of transcript capture we focused on morphologically defined regions with distinct marker genes, such as the choroid plexus and the retina (Figure 3D-F). Transcript capture was plotted on the H&E as a virtual *in situ*, with intensity relating to gene expression, resulting in a comparable stain with chromogenic *in situ* hybridisation (ISH) staining from the Allen Brain Atlas ^32,33^. Focusing on the fore-, mid-, and hindbrain regions, the expression of select marker genes showed high accordance with ISH (Figure S4C, F, I). For example, regionalized expression of the transcription factors *Neurod6* and *Pbx3* could be detected in the forebrain (Figure 3D) ^34^. As expected, the pan-neuronal marker Tubulin beta 3 (*Tubb3*) was absent in the choroid plexus, whilst being detected in the hindbrain (Figure 3E).

We proceeded to quantify the transcript density of transthyretin (*Ttr*) and atonal BHLH transcription factor 7 (*Atoh7*) across lines on the signal, observing a corresponding sharp rise and fall of signal intensity (Figure 3F, S3G). We find that the majority of these transcripts are captured in regions covered by tissue: the proportion of accumulated UMIs across the line profile that fall underneath tissue, measured as area under curve, yields values of 95/93% and 91/98% for *Ttr* and *Atoh7*, respectively. Transcripts observed in regions without tissue may be due to inaccuracies in manually defining the tissue boundary, or local lateral diffusion. The transition between a local minimum and maximum occurred within the length of a cell (between 7 and 11 µm) for *Ttr*. On the other hand, *Atoh7* exhibits a less sharp boundary (between 20 and 25 µm), which may be an effect of a wider region being sampled due to overall sparser and lower expression than *Ttr*.

While our localization analysis unveiled a predominantly local capture pattern, we also performed a crosstalk analysis to assess sources of spatial biases that may impact our interpretation (Sup. Note 1, Figure S3D). Focusing on ’blood’ and ’chondrocyte’ clusters, this analysis highlighted an increased gene marker mixing at pairs of cells within 20 µm proximity, underscoring the impact of local biases in clustering (Figure S3E). This consideration extends to challenges in segmentation and potential influences on dimensionality reduction and normalization approaches (Figure S3F).

### 5. Local capture of Open-ST reflects nuclear-cytoplasmic cell architecture

To assess the theoretical subcellular resolution of Open-ST based on the density of barcoded capture spots, we quantified the subcellular precision of transcript capture in data stemming from a coronal section of an adult mouse hippocampus hemisphere (Figure 3G).

Compared to other Open-ST datasets, this tissue exhibited lower UMI counts relative to the sequencing reads (Figure S3H, Table S2), likely due to an overall lower cell density resulting in a reduced initial RNA input across the capture area. The tissue’s variety of cell densities, however, rendered it suitable for assessing nuclear/cytoplasmic enrichment. Visualization of nuclear-retained *Malat1* and mitochondria-encoded transcript counts within the nuclear and cytoplasmic regions of the cell segmentation mask showed accurate transcript localization (Figure 3G). Average distribution of these transcripts in nuclear and cytoplasmic compartments was calculated for different cell densities across 1,791 regions (5,000 µm^2^ each) and projected onto a standardized cell (Figure 3H, S3I, Methods).

*Malat1* expression displayed significant enrichment in the nucleus (log_2_(Odds ratio cytoplasmic/nuclear) = -1.24, p_Fisher_<0.01), whereas mitochondrial transcripts were enriched in the cytoplasm (log_2_(Odds ratio cytoplasmic/nuclear) = 0.20, p_Fisher_<0.01). This was true across regions with variable cellular densities, whilst no significant enrichment was detected when off-setting the segmentation mask by 5 μm (Figure S3J). Importantly, when assessing the distribution of UMIs relative to their distance to the segmented nuclear edge, our observations of nuclear enrichment remained consistent (Figure S3K). The trends in UMI distribution around the nuclear periphery did not persist after a two-dimensional offset of the pairwise-aligned coordinates, further validating the localization patterns. In summary, we demonstrated that localized RNA capture on the high-density capture array reflected nuclear-cytoplasmic cell architecture.

### 6. Open-ST captures spatial cell type complexity in human primary tissues

To showcase the ability of Open-ST to dissect transcriptomic diversity in structurally complex tissues, we spatially sequenced a primary HNSCC tumor, and a healthy and metastatic lymph node from a single patient.

Annotation of unbiased clusters was performed based on literature-informed marker genes using a single-cell atlas reference ^24^. Additionally, cell type labels were refined by jointly clustering the lymph node tissues of the metastatic and healthy samples (Methods). We were able to identify distinct cell type signatures corresponding to the transcriptional diversity of tumor, stroma, and immune populations in the primary HNSCC (Figure 4A), healthy lymph node (Figure 4C), and metastatic lymph node (Figure 4D) datasets. As expected, the median number of genes and UMIs per cell in the tumor-enriched primary sample was higher than in the healthy lymph node (Figure 4B). The annotation of segmented cells matched the abundance of marker genes in space (white arrowheads, Figure 4E-G). Importantly, the major populations identified through the spatial transcriptomics data were independently outlined in H&E images by a professional pathologist (Figure S5D, F, H). The spatial separation of lymphoid tissue and infiltrating tumor in the metastatic lymph node annotated by the pathologist was also reflected in the neighborhood enrichment of Open-ST clusters (Figure S5C, H).

**Figure 4.**
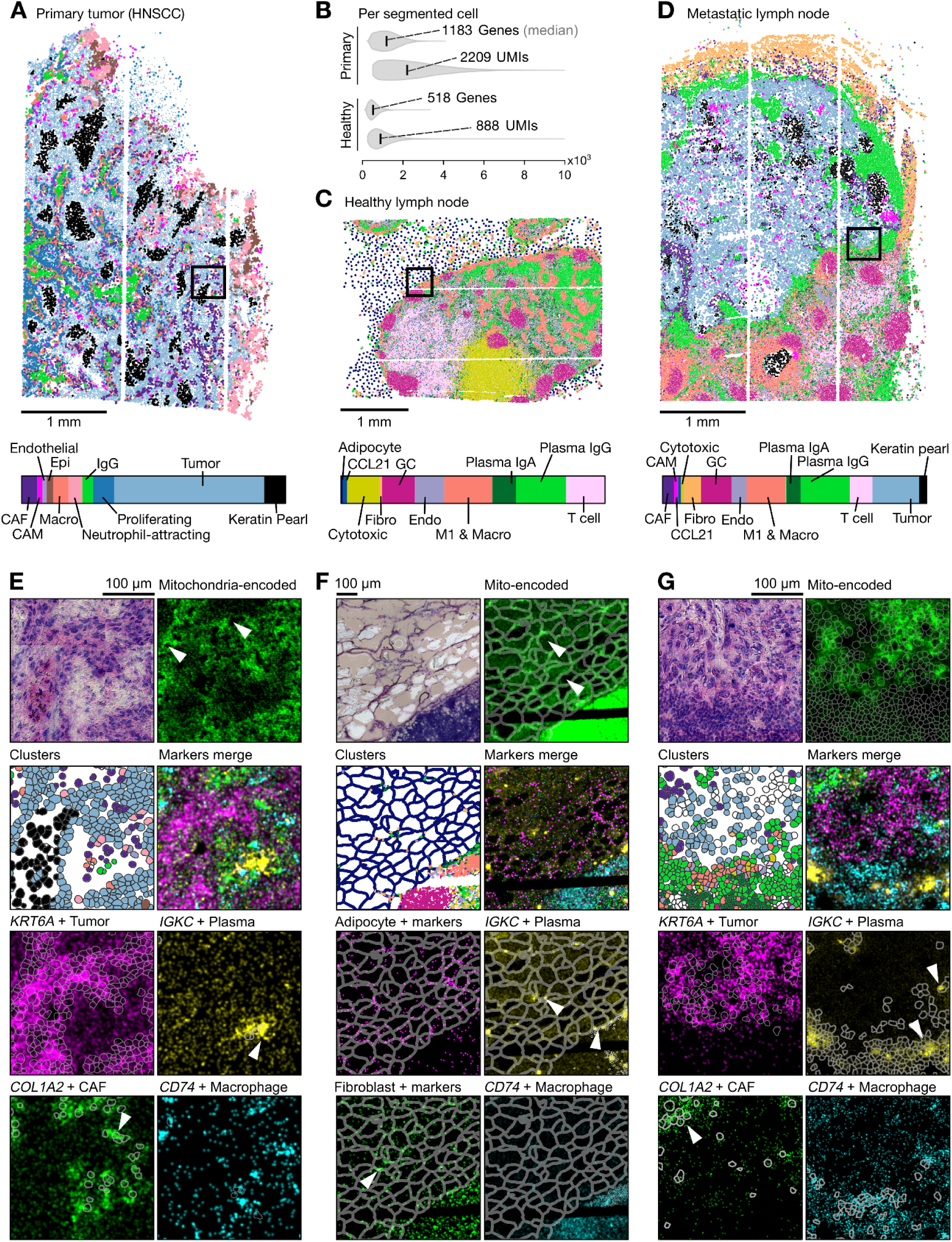
Open-ST accurately dissects cell type complexity in primary human tissues. **(A)** Spatial distribution of cell types (top) and cluster proportions (bottom) in a primary head and neck squamous cell carcinoma (HNSCC) section. **(B)** Distributions of the number of UMIs and genes per segmented cell in the primary HNSCC and healthy lymph node sections. **(C)** As in (A), but for the healthy lymph node. **(D)** As in (A), but for the metastatic lymph node. Spatial plots (A, C, D) visualize cell centers. CAF: cancer-associated fibroblast; CAM: cancer-associated macrophage; Epi: epithelial; IFNr: interferon production regulator; M1: M1 macrophage; Macro: macrophage; Fibro: fibroblast; Endo: endothelial; GC: germinal center. **(E-G)** Expression of selected marker genes in space for the primary HNSCC (E), the healthy lymph node (F) and the metastatic lymph node (G). Top left: H&E staining; top right: expression of mitochondria-encoded transcripts; second row: segmentation mask coloured by cluster (left) and gene expression of a marker from every cluster (right); third and fourth rows: gene expression of selected markers for different clusters, together with the corresponding segmentation masks. Pseudoimages show the expression of the indicated marker genes at spot resolution for the respective cell types (Methods). In (F) expression of all collagen genes is shown in “Fibroblasts and of *ADIPOQ, LEP, CIDEC, PLIN1, PLIN4, GHR, SORBS1, SIK2, SLC1A5, PDE3B, FABP4* in the “Adipocyte +” cluster. In the coloured segmentation masks, white cells represent those not passing counts or gene filters (Methods), or not assigned to any annotated cluster

The expression profile of the tumor cells reflects its epithelial origin; in fact, a high expression of keratins (*KRT6A, KRT5*), cell adhesion (*ITGB6)* and desmosomal *(DSG3)* markers is observed (Figure S5A, C; Table S4). Additionally, ‘keratin pearls’ (KP), which characterize squamous cell carcinomas, are found in both the primary and metastatic tumor tissues. These cells were found to colocalize with one another, in line with their known formation of distinct keratinized structures in the tumor bed (Figure S5A-C). These structures express the same epithelial markers found in the tumor cells, but also high levels of cornification-related molecules (*S100A10*) and desmosome markers (*DSP*), compatible with the activation of keratinocyte differentiation pathways. In the primary tumor, a cluster transcriptionally similar to the tumor and keratin pearl, was defined as ‘proliferating’; marker genes *RRM2*, *HMGB2*, and *CDCA5*, implicated in cell cycle regulation and/or DNA repair, were co-expressed in the section (Figure S5A, E). Furthermore, tumor cells, as well as cancer-associated macrophages (CAM), from both primary and metastatic tissues highly express the fatty acid binding protein-5 (*FABP5*), suggesting a dysregulation in the lipid metabolism of the cancer (Figure S5A, C).

Cancer-associated fibroblasts (CAF) (*COL1A1+*) were identified in the microenvironment of both tumor tissues, with a subset of CAFs in the metastatic tumor expressing high levels of *ACTA2*, suggesting a myofibroblast phenotype ^35^. Endothelial cells expressing *PECAM1* and *ACTA2* were identified in the primary tumor; while endothelial cells of the lymph nodes expressed *ACKR1*, a marker for post-capillary venules, and *CCL21*, a chemokine expressed in lymphatic endothelial cells ^36^.

Active macrophages expressing lysozyme (*LYZ*) were detected in all tissues. Moreover, pro-inflammatory M1-macrophages were identified in the metastatic lymph node, with high *TIMP1* expression in a subset of the cells. Cancer-associated macrophages (CAM) exhibited strong secreted phosphoprotein 1 (*SPP1)* expression, a gene implicated in macrophage polarity and previously associated with negative human papillomavirus (HPV) status and poor prognosis in HNSCC ^37^. This cluster was found to frequently colocalize with tumor cells, highlighting their interactive potential in the tissue microenvironment (TME). *SPP1*^+^ CAMs were previously identified to be enriched in areas of hypoxia in HNSCC, with *in vitro* studies indicating that hypoxia promotes *SPP1* expression ^38^. A cluster of neutrophil-attracting cells highly expressing *CXCL8* localize at the boundary of the primary tumor section and around the necrotic areas visible in the metastatic lymph node H&E staining (Figure S5A, E, I) ^39^. The *CCL21* marker gene is expressed by lymphoid endothelial cells in the T-cell zone and expressed also by T-cells (along with *CD3E*), and in interferon-responsive cytotoxic T cells (together with *IRF8* and *CD3E* expression). Within the lymph nodes, we detected multiple germinal centers, the microanatomical site of plasma and B cell formation. The germinal centers exhibited localized expression of *FDSCP*, a protein secreted by follicular dendritic cells, and immunoglobulin heavy constant mu (*IGHM*), suggesting an early germinal center status (Figure 4C, E; Figure S5B, C, G, H) ^40,41^. Furthermore, we classified two clusters with strong *IGHG3* or *IGHA1* expression as ‘IgG’ and ‘IgA’ plasma cells, respectively.

White adipose tissue was visibly surrounding the healthy lymph node in the annotated H&E images (Figure S5F). To retain the transcriptomic information of the uniocular adipocytes (i.e. containing a single lipid droplet in their cytoplasm), as well as that of the smaller cell populations of the adipose tissue stromal vascular fraction, our segmentation strategy was adapted in this sample (Methods). We unbiasedly identified cell type markers that enabled us to identify immune (e.g., *IGKC-*positive plasma cells) and stromal cells (e.g., collagen-expressing fibroblasts) within the adipocyte population (*FABP4, ADIPOQ, PLIN1, SORBS1, CIDEC* and *SIK2*), as pointed out by the white arrowheads in the pseudoimages (Methods) showing their localized marker expression (Figure 4F, S5B, G) ^42,43^.

Taken together, the above demonstrates that Open-ST accurately dissects the complex primary human tissue composition.

### 7. Open-ST reveals spatially-constrained heterogeneity in primary and metastatic HNSCC

Building on our exploration of the coarse cell types in the primary HNSCC and metastatic lymph node samples, we delved into the transcriptomic heterogeneity of tumor cells. By integrating and clustering 42,132 cells from both samples, previously annotated as ‘tumor’, ‘keratin pearl’ or ‘proliferating’, we identified 10 distinct transcriptomic states (T1-10), providing a detailed map of their heterogeneity, spatial distribution and stromal interactions (Figure 5 and S6) (Methods).

**Figure 5.**
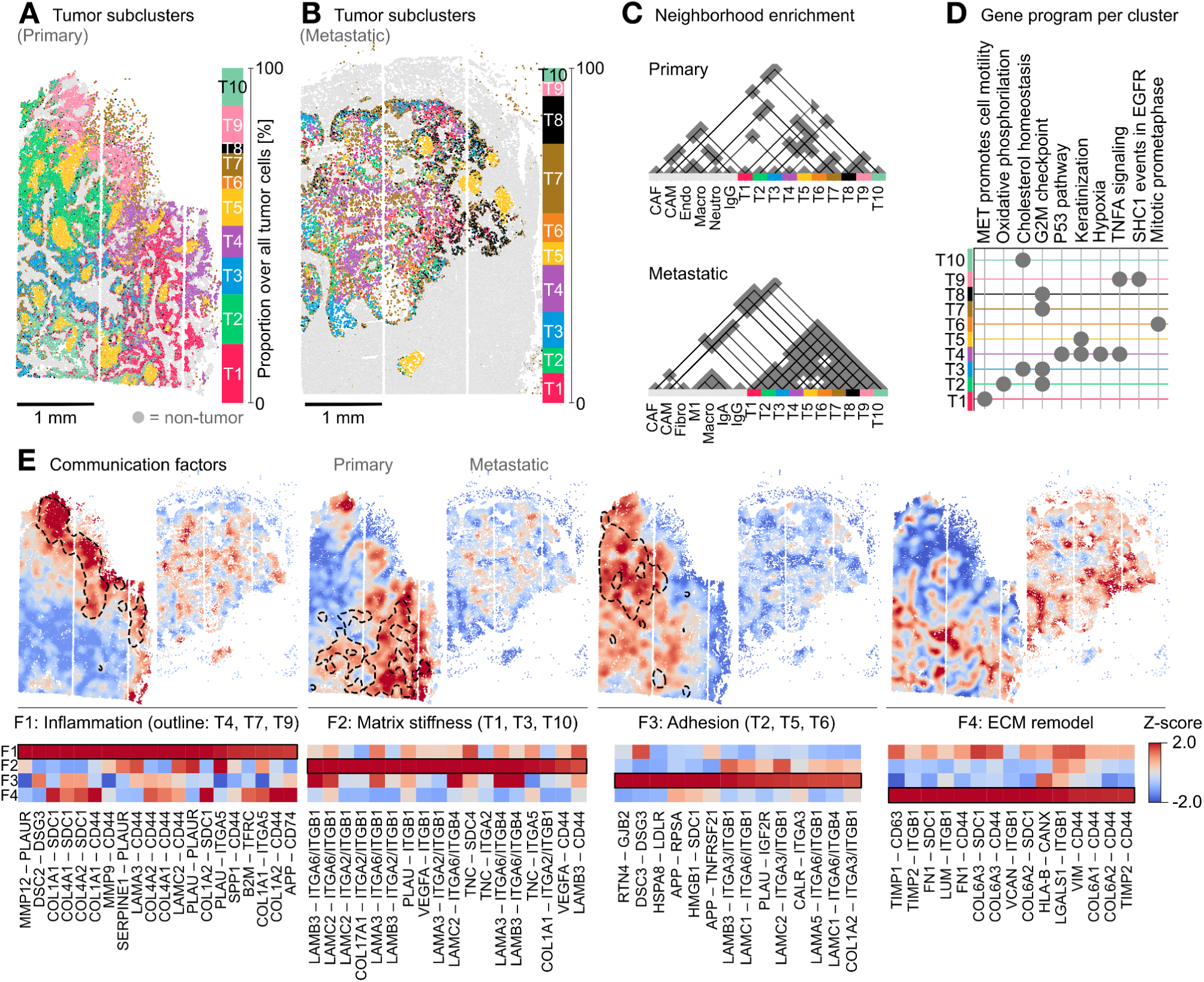
Transcriptomic tumor heterogeneity is organized into spatial domains with distinct communication signatures. **(A)** Spatial and transcriptomic heterogeneity of tumor cells across the primary tumor tissue. **(B)** As in (A), but in the metastatic tissue. **(C)** Spatial neighborhood enrichment of tumor and stroma populations. Significant spatial interactions (permutation test p-value < 0.05) between cells belonging to a pair of clusters are depicted by gray triangles, and connected by solid lines (Methods). **(D)** Normalized enrichment score (NES) in the tumor subclusters for the primary tumor tissue highlights differentially active gene programs. Dots are shown for cases where NES > 1 and FDR-adjusted p-value < 0.05 (Methods). **(E-F)** Cell-cell communication is organized as spatial motifs discovered with non-negative matrix factorization, that correlate with the tumor type heterogeneity in the primary (E) and metastatic (F) samples. Top: spatial visualization of factor contributions after z-normalization; bottom: min-max normalized communication score for top representative ligand-receptor pairs for each factor (Methods).

**Figure 6:**
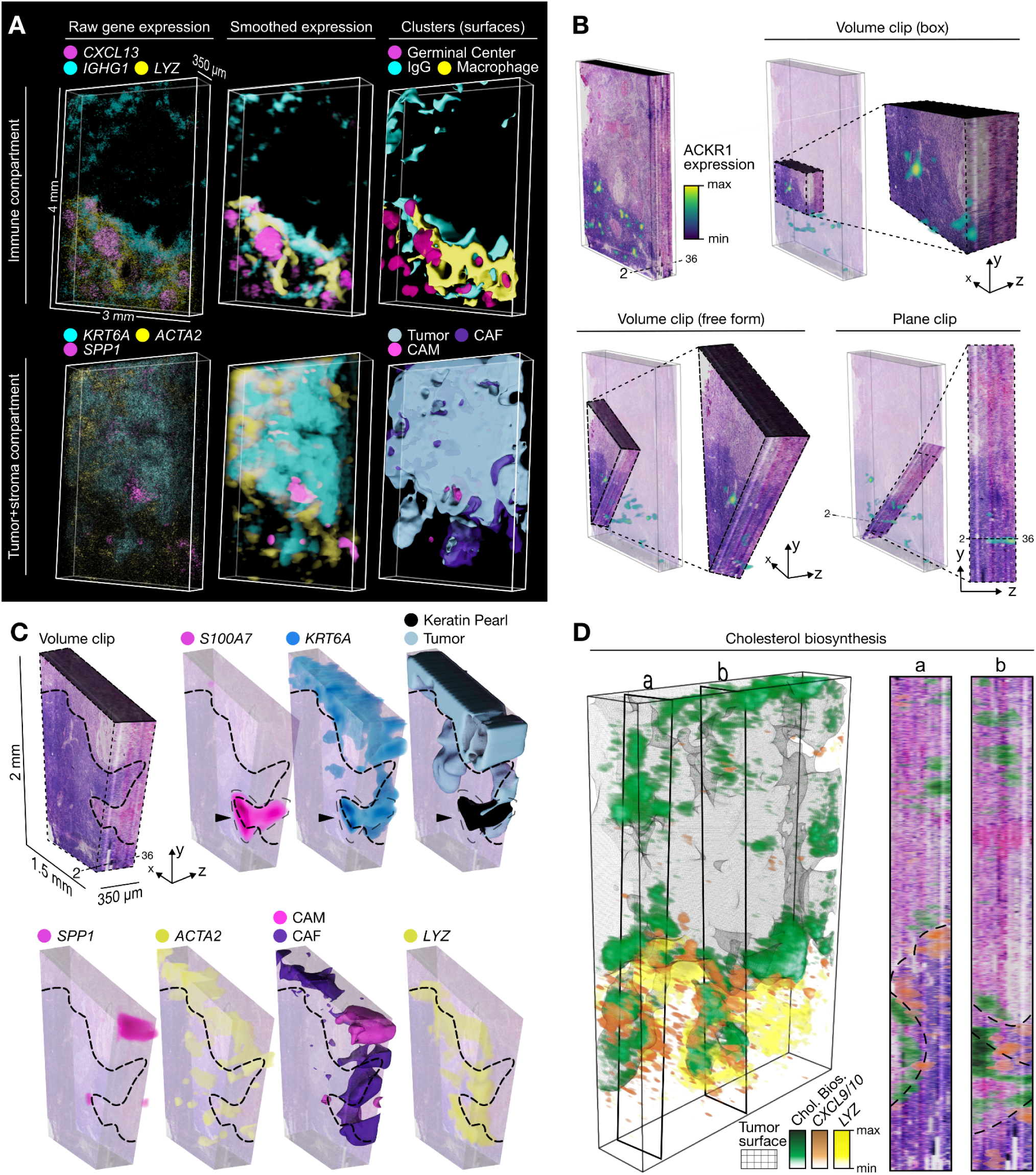
3D virtual tissue block: the metastatic lymph node. **(A)** 3D-visualizations highlight the spatial compartmentalization and continuity of gene expression. From left to right: gene expression levels of the markers rendered at each cell, smoothed expression, and 3D-rendering of clusters as smooth surfaces. Top: immune compartment represented by the germinal center (*CXCL13),* IgG plasma cells (*IGHG1*) and macrophages (*LYZ*); bottom: tumor and stroma compartments represented by epithelial cells (*KRT6A*), CAMs (*SPP1*) and CAFs (*ACTA2*). **(B)** Spatial expression of venular endothelial marker *ACKR1*. The virtual tissue block obstructs the view of expression in 3D. Rendering techniques allow exploration of the continuity of gene expression in 3D on the example of *ACKR1*. From left to right: volume clips (box or free-form) and plane clipping showing expression across the z-axis. Reduced opacity of the virtual tissue block surrounding the clip reveals *ACKR1* expression within tissue. **(C)** From left to right, a volume clip of the H&E tissue block shows the continuity of a keratin pearl structure that appears ‘disconnected’ from the main tumor mass in early sections. Gene expression supports this continuity along the z-axis, reflected by *S100A7* (specific to keratin pearls) and *KRT6A* (all tumor cells) over the semi-transparent tissue block. The stroma, demarcated by the expression of *SPP1* (CAMs) and *ACTA2* (CAFs), also follows this tumor structure penetrating the lymph node, whose boundary is rich in *LYZ, CXCL9* and *CXCL10* (pro-inflammatory macrophages). Dashed lines indicate the tumor/immune boundary as determined from the H&E. **(D)** Spatial expression of *LYZ* and the chemokines *CXCL9*/*CXCL10*, and spatial activity of the cholesterol biosynthesis gene sets from Reactome, quantified per segmented cell with AUCell and visualized as smoothed volumetric renderings (left) ^44^. Tumor surface is shown as a skeletal representation (wireframe) in gray. Gene expression and program activity are additionally shown in two cross sections (right, a-b).

The tumor transcriptomic states showed a distinct spatial organization across the primary and metastatic samples. In particular, tumor states in the primary tumor were organized into continuous spatial domains prevalent for a specific state (Figure 5A). In contrast, in the metastatic tissue tumor states appear more intermixed without clearly defined boundaries (Figure 5B). This observation was validated via neighborhood enrichment analysis (Figure 5C, Methods).

We further assessed the transcriptomic heterogeneity of these spatially-organized tumor states via differential gene expression (DE) and gene-set enrichment analysis (GSEA) (Methods). We detected state-enriched gene programs, with non-random spatial organization across the two tissues (Figure 5D, S6B-E, Methods). For instance, in the metastatic lymph node, we detected increased *cell cycle* activity at the boundary between tumor cells and lymphoid tissue (T2, T3, T6, T8), as well as around an hypoxic region (T4, T7). Notably, the *cholesterol biosynthesis* program (via *HMGCR*, *HMGCS1*, *DHCR7* and *DHCR24*) was spatially restricted to the tumor side of the tumor-lymphoid boundary (Figure S6E, F, Methods). On the other hand, the lymphoid (paratumor) side of the boundary was enriched in cells expressing *GZMA*, *GZMB*, *LYZ*, *CXCL9*, *CXCL10* and other markers of immune activation (Figure S6G).

To characterize the interplay between communication and transcriptomic identity, we performed receptor-ligand interaction analysis based on a consensus database (Methods), followed by spatially-aware factorization of communication scores to reveal common patterns of communication. This factorization revealed four non-negative factors across primary tumor and metastasis (Figure 5E, F, Methods), each with unique communication signatures, which we termed based on literature-informed annotations of their main ligand-receptor pairs: inflammation (F1), matrix stiffness (F2), adhesion (F3) and ECM remodeling (F4). These factors displayed a higher degree of spatial organization in the primary tumor compared to the metastatic, following a more intermixed organization of tumor states.

We found different tumor states to be restricted to specific cell-cell interaction hotspots. In the primary tissue, T1 was found in a CAF-rich stroma and showed cell motility phenotype, hinted by the upregulation of *LAMC2, PTHLH* and *ITGB1* (Figure S6A), as well as spatially-restricted activity of communication axes related to matrix stiffness and migration, such as integrin or syndecan-mediated communication (Figure 5E), similar to previously reported partial EMT subcluster of HNSCC tumor cells with upregulation of *LAMC2, TGFBI* and *SNAI2* ^24^. This set of tumor cells was the only one with a spatial enrichment with CAFs in the primary tumor, which subsequently surrounded endothelial cells. T2 showed a predominant proliferation phenotype. It was enriched within a communication hotspot for cell adhesion mediated via desmoglein-desmocollin interactions, in the primary tumor only showing spatial interactions with tumor cells (T6, T8). Tumor states T3 also had marked proliferation signatures, additionally showing metabolic phenotypes, particularly cholesterol biosynthesis. It localized to the communication hotspot for matrix stiffness, surrounded by a microenvironment only composed of other tumor cells, remarkably T6 with similar proliferation phenotype, which neighbored macrophages. T4 and T9 showed phenotypes involving TNF-α signaling, hypoxia and other stress-induced pathways and localized near receptor-ligand interaction hotspots involving metalloproteases (*MMP9-CD44*, *MMP12-PLAUR*) and osteopontin (*SPP1*-*CD44*), suggesting inflammatory response (Figure 5D, E; S6D). Moreover, T4 and T9 were in close proximity to neutrophil-attracting, CAM populations (Figure 5C), and T7 localized to necrotic regions visible in the H&E (Figure S5H). T5, in agreement with the coarse clustering, displayed significant keratinization signatures and activity of the desmosome pathway, a known signaling mode for cell adhesion. T8 had low representation and sparse spatial localization in the primary tissue, and was more prevalent in the metastasis. T10 was abundant and spatially-organized in the primary tissue, in close proximity to cornified cells (T5) as well as CAMs, and showed a cholesterol biosynthesis phenotype similar to T3. Lastly, we globally detected differentially expressed genes between the two samples across all tumor cells: among other hits, a strikingly strong *AMTN* expression was identified exclusively in the metastatic lymph node, while *FGFBP1* and *PRR9* were exclusively detected in the primary tissue (Supplementary Table S7) (Methods).

In summary, Open-ST reveals diverse tumor cell states coexisting in close proximity within the primary tumor, in spatially-restricted domains whose organization closely follows that of cell-cell communication hotspots.

### 8. Exploring 3D virtual tissue blocks: cell types, gene programs, and receptor-ligand interactions in 3D

Open-ST software enables the interrogation of gene expression in 3D via volumetric renderings, and provides smoothed representations for enhanced clarity (Methods). As an example, we visualize the lymphoid, metastatic tumor and stroma tissue compartments via the expression of selected marker genes in 3D (Figure 6A).

These representations of gene expression coincide with the surface renderings generated from transcriptomic clusters, obtained after transferring the annotations from section 4 to the remaining 18 Open-ST sections (Methods). In particular, we classified the over 1 million cells into 14 major types, spanning tumor, immune, and stromal compartments. The spatial distributions and proportions of cell types in 3D were consistent across consecutive sections (Figure S7A-B). Our transcriptome annotation was also in agreement with the manual pathologist annotation of section 36 (Figure S7C). Furthermore, gene expression levels within section 4 were highly correlated with those of the most distant section 36, displaying reproducible RNA capture throughout the tissue block (Figure S7D).

Due to the high reproducibility across sections, our reconstructed tissue can be utilized for comprehensive exploratory data analysis, querying both the transcriptomic and imaging modalities from any perspective in 3D space (Methods). This allows exploration of gene expression in any direction regardless of the sectioning plane by leveraging rendering techniques such as volume cropping and plane clipping (Figure 6B). To illustrate Open-ST’s capabilities, we visualized the spatial gene expression of *ACKR1*, an endothelial marker restricted to post-capillary venules ^45,46,47^. Querying the tissue with a plane clip, which is equivalent to cutting the tissue using a different sectioning plane, allowed us to follow a post-capillary venule across the z-dimension demarcated by *ACKR1* and reveals how its expression is confined to the immune regions of the metastatic lymph node, forming a network-like structure (Figure 6B, bottom right).

To further showcase the capabilities of 3D reconstruction, we examined the keratin pearl which appeared as an isolated structure within the lymph node region in the initial sections (Figure 4D). Upon closer inspection, our 3D reconstruction revealed its connection to the metastatic mass in the later sections, demonstrating continuity across imaging and transcriptomic modalities (Figure 6C). Similarly, we could follow the continuity of the invading tumor and its stroma within the lymph node context via the markers *SPP1,* which marks CAMs, *ACTA2,* which highlights CAFs surrounding the tumor and is proximal to lymph node populations, and *LYZ,* which marks macrophages and is a transcriptomic feature located at the lymph sinus, as well on invaded tissue ^48^. This structure is in general found in close proximity to afferent lymphatic vessels, and sinus macrophages are the first to be exposed to the metastasis ^49^.

The power of 3D reconstruction extends well beyond the study of cell types and tissue structures. To further demonstrate its potential, we inspected gene program activities in 3D. Specifically, we validated the three-dimensional organization of cells with high cholesterol biosynthesis gene set activity at the boundary between tumor and lymphoid tissue, in close proximity to the sinus, hinted by the presence of *LYZ*+, *CXCL9*+ and *CXCL10+* cells (Figure 6D, Methods). 3D visualization further revealed the spatial activity of the G2M checkpoint, additionally around areas with hypoxic phenotype precisely located at the core of the tumor mass (Figure S7F), as previously hinted in a single 2D section (Figure S6E).

In summary, Open-ST provides a versatile and powerful framework for comprehensive analysis of gene expression in 2D and 3D, including the unveiling of molecular and/or cellular spatial structures obscured in traditional 2D representations. We anticipate Open-ST to accelerate the identification of spatial molecular mechanisms in 2D and 3D.

## Discussion

Here, we introduced Open-ST, a framework of experimental and computational tools for the efficient capture and interrogation of transcripts in tissue space at subcellular resolution.

We illustrated the quality of our data in the stereotypical developing E13 mouse head, with the transcript localization showing high accordance with *in situ* hybridizations from the Allen Developing Mouse Brain Atlas. We further demonstrated the wide applicability of Open-ST by spatially sequencing structurally complex primary human tissues of clinical interest. Benchmarked against comparable spatial technologies, Open-ST outperforms in terms of UMI counts per cell at similar sequencing depth, while retaining a low reads-to-UMIs ratio.

Open-ST allowed us to recapitulate tumor heterogeneity at single-cell resolution, from primary and metastatic lymph node samples of the same patient. In the primary tissue, we discovered heterogeneous states of tumor and stromal cells, restricted to specific spatial domains with enrichment of specific gene programs and receptor-ligand interaction hotspots.

These states involved inflammatory response, epithelial differentiation and keratinization, proliferation and matrix remodeling with migratory potential. We hypothesize that this last population, situated in close proximity to CAF/endothelial-rich stroma, serves as the origin of metastatic cells. Tumor cell and stromal heterogeneity were maintained in the metastatic tissue suggesting co-migration during metastasis, a phenomenon previously suggested in single-cell studies ^24,50^. The differences in spatial arrangement of tumor subclusters in the metastasis suggest that transcriptomic states are established around specific communication hotspots in the primary tissue, and are mostly maintained upon new microenvironments. While transcriptomic states were generally similar for the detected subclusters, we found unique features in the metastatic tissue, such as the upregulation of *AMTN*, potentially caused by the differences in signaling at the new microenvironment ^51,52^. Moreover, 3D visualization confirmed the spatial patterns of gene program activity in the metastatic lymph node. At the tumor/lymphoid boundary, we observed the upregulation of a cholesterol biosynthesis signature, and gene expression patterns (i.e., *LYZ*+ *CXCL9*+ *CXCL10*+ macrophages) that hint the structural integrity of the subcapsular sinus, a structure located at the outer layer of the lymph node. We hypothesize that the growth of the tumor mass along the boundary pushes the lymphoid tissue rather than breaking it. This provides mechanical cues, linked to the regulation of lipid metabolism ^53,54^. The upregulation of cholesterol metabolism is in itself an immune-evasive mechanism: tumor cells with cholesterol-enriched membranes have been shown to impair T-cell mediated cytotoxicity ^55^. Moreover, previous studies report dysregulation of cholesterol metabolism in HNSCC as a potential therapeutic target, in particular, the enzyme HMGCR (HMG-CoA reductase) ^56,57^. Similarly, a fatty acid metabolism signature at the tumor/paratumor boundary has been reported in liver cancer ^4^.

All our analyses were obtained by using the Open-ST computational toolkit for the diverse datasets showcased in this study. We anticipate that this toolkit can be applied to other experimental setups and processed samples, requiring minimal user intervention. Additionally, the modular approach enables the use of alternative tools and algorithms for any (pre)processing step, when necessary. Altogether, we provide detailed experimental resources and open-source computational tools that hold the potential to render Open-ST as a standard for community-driven generation and analysis of spatial transcriptomics.

### Limitations and future developments

Open-ST is currently limited to the study of polyadenylated transcripts and carries a 3’ capture bias. The library preparation can be adapted to retain the entire length of the transcripts, enabling the spatial analysis of isoform diversity. Similarly, the capture oligos can be modified and designed to capture specific targets, such as specific splice variants or bacterial 16S transcripts, allowing the phylogenetic classification of microbes. Moreover, we anticipate to render Open-ST compatible with formalin-fixed paraffin embedded tissues, widening its applicability. Finally, Open-ST requires an initial financial investment to generate the capture areas, and is therefore not well-suited for low-throughput experiments. Collaborations or centralized capture area generation by institutes can overcome this hurdle.

Cell segmentation is currently guided by nuclear staining only, limiting its precision and contributing to the cross-contamination of cellular transcriptomes. This can be addressed by introducing membrane staining, additionally enabling the detection of multinucleated cells. Incorporating immunofluorescence or immunohistochemical stainings may negatively affect RNA quality and/or localization of RNA capture, and thus, require separate optimization. The discrepancy in z-versus x-y resolution (10 μm vs 0.6 μm) presents an additional limitation, as transcripts deriving from different cells layered along the z-dimension may be captured, confounding single-cell level analyses. Future development of computational methods will allow deconvolution of this data.

Currently, no standardized data analysis pipeline exists to account for artifactual local biases of sequencing-based spatial transcriptomics methods such as Open-ST: anisotropic lateral diffusion, spatially autocorrelated background signal, and misassigned transcripts due to inaccurate two-dimensional segmentation or the lack of resolution in the z-axis. Here, we leveraged standard workflows tailored for scRNA-seq that rely on normalization and manifold learning from the cells-by-genes matrix, without considering spatial autocorrelation as a source of covariance ^58–60^. We anticipate that spatial autocorrelation might break the assumptions underlying normalization and manifold learning, leading to false discoveries, such as spurious cell types, and inaccurate differential gene expression and activity of programs. We underscore the importance of future developments in spatially-aware normalization, dimensionality reduction and clustering, to address questions that rely on modeling count data in space, such as differential expression analysis, as well as analysis relying on manifold structure, such as cell typing, pseudotime and RNA velocity.

## Acknowledgements

We thank all members of the Rajewsky lab for critical and helpful discussions. We thank Alexandra Tschernycheff, Veronica Jakobi and Margareta Herzog for supporting the organization of the lab. Thank you to Cledi Alicia Cerda-Jara and Gwendolin Thomas for providing mouse samples and Lieke van de Haar for discussions on mouse brain anatomy. We also thank Salah Ayoub, Marvin Jens and our former lab member Asija Diag for their input to the spatial transcriptomics method development. We thank the Genomics Technology Platform at the MDC/BIH@Charité, particularly Tatiana Borodina and Jeannine Wilde, for performing sequencing runs and providing us with custom flow cells.

## Illustrations in Figures 1D and 2A and D were created with Biorender

M.S. was financially supported by the Berlin Institute of Health at Charité (BIH) between December 2020 and February 2022, D.L.-P. by the Helmholtz Einstein International Berlin Research School in Data Science (HEIBRiDS) program of the Helmholtz Association and E.S. received an Erasmus fellowship. L.S. received financial support from the International Research Training Group (IRTG) 2403 program of the Humboldt-Universität zu Berlin. T.M.P. was financially supported by the Berlin School of Integrative Oncology through the GSSP program of the German Academy of Exchange Service (DAAD) and by the Add-on Fellowship of the Joachim Herz Foundation. J.A received funding from the MDC/NYU PhD Exchange Program, S.S.T from the Berlin Institute of Health at Charité (BIH) between December 2020 and July 2022, and A.B from the Leibnizpreis DFG RA 838/5-1. N.K. was financially supported by the Deutsche Forschungsgemeinschaft (DFG) grant number KA 5006/1-1. G.M received financial support via a BIH Visiting Professorship “Single Cell Genetics and Epigenetics of Patient Derived Tumor and Brain Organoids” by the Stiftung Charité from 2018 to 2022. N.R. thanks: the Leibniz prize of the Deutsche Forschungsgemeinschaft (DFG) (grant number RA 838/5-1) , the DFG Exzellenzcluster 2049 NeuroCure, the Deutsches Zentrum für Herz-Kreislauf-Forschung (DZHK) grant numbers 81Z0100105 and 81X2100155, and the Chan Zuckerberg Foundation (CZI)/Seed Network.

## Author contributions

G.M. and N.R. conceived the project. J.A., S.A., A.B., S.E., J.L., G.M., N.R., S.S.T., M.S., and E.S. contributed to the experimental method development. N.K., G.M., D.L.-P., T.M.P., N.R., M.S., and E.S. designed the experiments. S.D. and S.S. collected the human samples and performed the pathological analysis. M.S. and E.S. processed all samples except for the mouse head, which was processed jointly with J.L.. N.K. and D.L.-P. performed the computational method development. D.L.-P led the computational analysis, supported by N.K. and L.S.. N.K., G.M., and N.R. supervised the project. S.A. and E.S. were supported by M.P. and E.F., respectively. N.K., D.L.-P., N.R., M.S., E.S., and L.S. wrote the original draft. All authors revised the manuscript and approved the final version.

## Methods

### Resource availability

#### Lead contact

Further information and requests for resources and reagents may be directed to Nikolaus Rajewsky.

#### Materials availability

The design of the 3D-printed cutting guide is provided in our online resource (https://rajewsky-lab.github.io/openst) . All other materials used are commercially available.

#### Data and code availability

Open-ST RNA-seq and microscopy raw and processed data have been deposited at GEO and will be publicly available as of the date of peer-reviewed publication.

All original code has been deposited at https://github.com/rajewsky-lab/openst (accessed 3 November 2023) and is publicly available as of the date of publication. DOIs are listed in the key resources table.

Any additional information required to reanalyze the data reported in this paper is available from the lead contact upon request.

### Experimental model and subject details

#### Mouse tissue sample

Mouse adult brain was collected from a postnatal-day 60 C57BL/6N wild-type male mouse (RRID:MGI:2159965). Embryonic stage 13 (E13) mouse head was collected from a C57BL/6N wild-type mouse (RRID:MGI:2159965). Tissue was immediately embedded in Tissue-Tek® O.C.T.™ Compound (Sakura, 4583) on powdered dry ice and stored at -80°C. The left hemisphere of the adult brain was embedded and sectioned coronally. Mouse E13 head was embedded and sectioned sagittally.

Animal care and mouse work were conducted according to the guidelines of the Institutional Animal Care and Use Committee of the Max Delbrück Center for Molecular Medicine and the Landesamt für Gesundheit und Soziales of the federal state of Berlin (Berlin, Germany).

#### Human patient characteristics and sample collection

The resected head and neck tumor, as well as a normal and a metastatic lymph node, were collected from a 56 year-old male patient with a moderately differentiated, keratinizing squamous cell carcinoma (SCC) in the larynx.

Staging workup by cranial computed tomography (CCT) and Positron emission tomography–computed tomography (PET-CT) showed bilateral cervical lymph node metastases, but no distant metastases. A laryngectomy and bilateral neck dissection was performed and the patient received an ambulant adjuvant radiotherapy.

The pathological examination revealed tumor infiltration of the supraglottis, glottis, subglottis regions, pre-epiglottic soft tissue, and two cervical lymph node metastases with a maximum diameter of 12 mm and without extracapsular invasion.

A part of the primary tumor, as well as half of resected level III cervical lymph nodes with and without metastasis, were snap-frozen after surgery and embedded in pre-cooled Tissue-Tek® O.C.T.™ Compound (Sakura, 4583) on powdered dry ice. Tissue was stored at -80°C before and after embedding and used for Open-ST.

The remaining resected specimen was formalin-fixed, decalcified, cut and paraffin-embedded (FFPE) for histological examination. The pathological tumor classification was as follows: pT3 pN2c (2/29) L0 V0 Pn0 G2 R0 (according to the 8th edition of the TNM classification (AJCC)). The FFPE tissue was stored at room temperature at the archive of the Institute of Pathology at the Charité University Hospital, Campus Mitte. The study was performed according to the ethical principles for medical research of the Declaration of Helsinki and approval was approved by the Ethics Committee of the Charité University Medical Department in Berlin (EA4/082/22).

### Method details

#### Capture array generation and disassembly

The capture array was generated as in Cho *et al.* (2021) with several protocol adaptations ^16^. The synthetic HDMI32-DraI library was produced using the Ultramer service from IDT. The library was sequenced on an Illumina® NovaSeq 6000 S4 flow cell (35 cycles), at a loading concentration of 200 pM. A single-end 37 cycle read was sequenced, using Read1-DraI (IDT) as a custom primer. A custom sequencing recipe was used, stopping the run immediately after read 1 prior to on-instrument washes (Data S1).

After sequencing, enzymatic reactions were performed by pipetting reaction mixes into the flow cell lanes, covering the inlets/outlets with tape for the incubations. First, the flow cell was washed by flowing through 500 uL ultrapure water, then incubated overnight with DraI mix (2 U/uL DraI enzyme (NEB, R0129) in 1X CutSmart buffer) at 37°C. A repeat 5h incubation at 37°C with fresh mix was done, to ensure efficient digestion in all areas. The DraI cuts the double stranded DNA of the clusters after the poly-d(T) tail. After washes with 80% ethanol, then ultrapure water, the flow cell was incubated with Exonuclease I mix (1 U/uL Exo I (NEB, M0293L) in 1x Exo I buffer) for 45 min at 37°C. Subsequently, the flow cell was washed three times with ultrapure water before separating the two glass surfaces by scoring along the sides with a scalpel.

The second strand was denatured by washing the opened flow cell in a beaker of 0.1 N NaOH for 5 min. After denaturation three washes each with 0.1M Tris HCl (pH 7.5) and ultrapure water were performed.

Using a 3D-printed tool and a glass cutter (available at https://rajewsky-lab.github.io/openst), scores were made along the flow cell top or bottom pieces at the desired breaking points. The cutting tool allows clean scores on the surface without oligos, without damaging the capture area surface. By applying even pressure at the scored sites, the flow cell was broken into approximately 3x4 mm capture area pieces. The first and last 1.5 cm of the S4 flow cell was removed, as these areas are not images by the sequencer and thus do not contain registered barcodes. The cutting guide was printed with 1.75 mm polylactic acid (PLA) filament on the Pro 3 Dual Extruder 3D printer from Raise 3D. The *.stl* file was prepared for printing using IdeaMaker v.4.3.2 as the 3D slicing software. Capture areas from two different flow cells were used for experiments: fc_1 (human samples, embryonic mouse brain) and fc_2 (mouse hippocampus).

#### RNA quality control

To assess RNA quality, total RNA was extracted from cryosections lysed in Trizol using the Direct-zol RNA Miniprep kit (Zymo Research, R2050) according to the manufacturer’s instructions. Concentrations were measured using the Nanodrop-1000 spectrophotometer (Thermo Fisher) and RIN values assessed using the Agilent 2200 TapeStation (High sensitivity RNA kit, Agilent).

#### Cryosectioning and fixation

OCT-embedded fresh frozen tissue was sectioned at 10 um thickness using a CryoStar NX70 cryostat (Epredia^TM^). Tissue sections were placed and melted onto the capture area and fixed in methanol at -20°C for 30 min.

#### H&E staining and imaging

The section was dried before H&E staining by 1 min incubation with isopropanol and air-drying at room temperature. For H&E staining, Mayer’s haematoxylin (Agilent, S3309) was applied for 5 min, the section was washed ten times in water and incubated with bluing buffer (Agilent, CS702) for 2 min. After washing in dH20, the tissue was treated with a 1:1 dilution of eosin Y (Sigma, HT110216) and 0.45 M tris-acetic acid pH 6 for 1 min. The sections were washed in water and left to air-dry completely before imaging. The tissues were imaged in brightfield with a 20x objective on the Keyence BZ-X710 inverted fluorescence phase contrast microscope, placed dry onto a # 1.5 coverslip.

#### Permeabilization time optimization

To select a suitable permeabilization condition we firstly tried the tissue optimization assay ^61^. Fluorescently labeled nucleotides are integrated into the cDNA during RNA retrotranscription; the signal intensity should correspond to the amount of RNA captured, with localization of signal indicating possible diffusion.

In our experience this assay was not reliable to quantify capture efficiency, so instead we define RNA capture efficiency using qPCR, where low amplification cycling numbers correspond to a higher concentration of starting material.

We tested different concentrations of pepsin (0.7-1.4 U/uL) for different incubation times (0, 15, 30, 45, 60 min) at 37°C. Library preparation was done as for a regular Open-ST assay, but only until second strand synthesis. A qPCR was performed as described in the methods section “PCR cycling number assessment”. If multiple samples amplified earliest together, we chose the shorter time or lower pepsin concentration for permeabilization.

#### mRNA release and capture

Reactions on the capture array were performed in a 16-well ProChamber microarray gasket (GraceBiolabs, 645508). Each flow-cell piece was placed in one gasket well with the tissue on the capture area facing up. To cover the entire surface a volume of 100 ul was used for all subsequently described reactions. mRNA was released by treating the tissue with pepsin (P7000, Sigma) in 2x SSC (pH 2.5) at 37°C, with time and concentration differing between the tissue. The primary human samples were permeabilized for 45 min with 1.4 U/uL pepsin. The mouse hippocampus and E13 head were permeabilized with 0.7 U/uL pepsin for 30 min.

The tissue was washed with a reverse transcription buffer (1x SuperScript IV buffer (Thermo Fisher), 1 U/uL Ribolock RNase inhibitor (Thermo Fisher)).

Next, an overnight incubation was done at 42°C with reverse transcription mix (6.67 U/uL SuperScript IV (ThermoFisher), 1x SuperScript IV buffer, 5 mM DTT 0.187 mg/mL BSA, 1 mM dNTPs, 1 U/uL Ribolock (ThermoFisher). The wells of the gaskets were sealed with strips of plate sealing tape to prevent evaporation.

#### Tissue removal

Tissue removal was done as in Cho *et al.* (2021), but with 100 uL mix or wash per sample to completely cover the flow cell piece ^16^. Tissue digestion was added after the reverse transcription reaction (100 mM Tris-HCl pH 8.0, 200 mM NaCl, 2% SDS, 5 mM EDTA, 16 mU/uL proteinase K (NEB, P8107S)). An incubation for 40 min at 37°C was followed by three ultrapure water washes, three 5 min 0.1 N NaOH incubations to denature the hybridized mRNA, three 0.1 M Tris (pH 7.5) washes for neutralization and finally three ultrapure water washes. The capture area was visually inspected to confirm tissue removal before proceeding with 2^nd^ strand synthesis.

#### Second strand synthesis and purification

Following the permeabilization step, the tissues were incubated with a Second Strand Synthesis mix for 2h at 37°C. The mix contains 1x NEBuffer-2 (NEB, #B7002S), 10 µM of Randomer (custom DNA oligo, IDT), 1 mM of dNTPs mix, 0.5 U/µl Klenow exo (-) Fragment (NEB, #M0212L). The random priming site serves as a UMI in downstream analysis.

The slides were washed three times with ultrapure water and incubated twice for 5 min with 100 uL 0.1 N NaOH to elute the strand. To neutralize the product Tris-HCl pH 7.5 was added to reach 0.125 M. AMPure XP magnetic beads (BeckmanCoulter, #A63881) were used to clean up and concentrate the second strand eluate to 82.5 μL, following the manufacturer’s instructions.

#### PCR cycling number assessment

qPCR was performed on the StepOne™ Real-Time PCR System (Applied Biosystems) to assess the number of amplification cycles needed (Figure S1F).

Around 3% of the purified Secondary Strand (2.5 μL) was used as input, in a mix of 1x Blue S’Green qPCR mix plus ROX (Biozym, #331416) and 1 μM p5 Fwd and p7 Rev indexing primers (Custom DNA oligo, IDT). Cycling conditions were as follows: 95°C for 2 min, followed by 40 cycles of 95°C for 5 sec and 60°C for 30 sec.

The threshold was set at 50% of the peak ΔRn. For each sample the cycle number at the intersection of the threshold and amplification curve was determined. Five cycles less were considered for the Secondary Strand amplification, to account for the input. For all the samples 12 or 13 cycles were selected for amplification.

#### Library construction

The libraries were prepared for amplification, combining 80 μL of purified second strand with a final 1x KAPA HiFi Library Amp kit (Roche, #07958960001), 1μM of p5 Fwd primer and p7 indexing Rev primer in a volume of 200 μL, split into 4 PCR tubes of 50 μL each. The following cycling conditions were used: initial denaturation for 3 min at 95°C, x cycles of 95°C for 30 sec, 60°C for 1 min and 72°C for 1 min, a 2 min final elongation at 72°C and hold at 4°C.

The AMPure XP magnetic beads (BeckmanCoulter, #A63881) were used at a 1:1 ratio to clean and concentrate the library into a final volume of 20 μl following the manufacturer’s instructions. To completely remove short artifacts, such as primer dimers, libraries were selected to have a length from 300-400 bp to 1100 bp, according to library composition and shape (Supp Fig 1E). Size selection was performed using 1.5% agarose gel cassettes on the BluePippin or PippinHT (Sage science, BDF1510 or HTC1510) following the manufacturer’s instructions. The product was quantified using the Qubit™ dsDNA HS Kit (Invitrogen, Q32851) and the Bioanalyzer Agilent High Sensitivity DNA Kit (Agilent, 5067-4626).

#### Spatial transcriptome sequencing

Libraries were quantified for sequencing using the KAPA Library Quantification Kit (optimized for Roche® LightCycler 480, 07960298001). Sequencing was performed on the Illumina® NovaSeq 6000 and NextSeq 2000 sequencing systems, with 130 pM and 650 pM loading, respectively, and 1-5% PhiX spike-in. A minimum of 28 cycles for read 1 and 90 cycles for read 2, as well as an 8-cycle index 1 read, were used.

#### Quality control and spatial reconstruction of barcode sequencing

The barcode sequencing for capture area generation enables the mapping of spatial barcodes to their corresponding spatial coordinates within the flow cell. Per flow cell, we processed each of the 3,744 tiles leveraging bcl2fastq v2.20.0.422. Subsequently, the barcode_preprocessing function from our openst tools was used to trim barcodes, compute their reverse complement ("cell_bc"), and supplement these with spatial coordinates ("xcoord" and "ycoord"), ultimately generating a single coordinate file per tile. Notably, the spatial coordinates acquired with bcl2fastq are in a tile-specific coordinate system. Consequently, mapping to a global coordinate system becomes necessary for samples spanning multiple tiles. This was done with the `puck_collection` functionality from spacemake v0.7.3, using the provided NovaSeq S4 coordinate system.

#### Preprocessing, mapping & gene quantification of spatial transcriptome sequencing

The transformation of raw sequencing data into spatially-mapped expression matrices was carried out utilizing spacemake ^27^, an automated pipeline designed for the preprocessing, alignment, and quantification of single-cell and spatial transcriptomics data.

#### Preprocessing

Raw basecalls were demultiplexed with bcl2fastq v2.20.0.422, yielding two FASTQ files per sample: the Read 1 (R1) file, containing the spatial barcode, and the Read 2 (R2) file, containing the unique molecular identifier (UMI) and transcriptomic data. The R1 and R2 FASTQ pairs were merged into a single bam file per sample. Finally, the consolidated bam files underwent trimming of p5 adapter sequences and polyA tails using *TrimStartingSequence* and *PolyATrimmer* from Drop-seq tools v2.5.1, respectively. In spacemake, this preprocessing strategy is represented by the “open-st” barcode_flavor and run_mode.

#### Mapping

Initially, preprocessed and trimmed reads were aligned to the PhiX reference sequence, with aligned reads subsequently discarded. Remaining reads were subjected to alignment against the corresponding rRNA reference, dependent on the sample’s origin (mouse or human). Aligned rRNA reads were discarded. These two steps were performed with bowtie2 v2.5.1 for fast alignment. The final step involved mapping the remaining unmapped reads to the species genome using STAR v2.7.10b, adhering to default parameters within spacemake v0.7.3. For mouse-origin reads, GRCm39 primary assembly with the Gencode vM30 annotation was utilized. Likewise, the GRCh38 genome reference with the Gencode v43 basic annotation was utilized for alignment of human-origin reads. The above mapping strategy can be reproduced in spacemake >=v0.7 by specifying a map_strategy as follows: “bowtie2:phiX->bowtie2:rRNA->STAR:genome:final”.

#### Quantification

Uniquely mapped reads were spatially matched against a library of spatial barcodes derived from the initial flow cell sequencing. Multi-mapping reads were discarded, except for the cases where a read mapped to one genic and one intergenic locus only; in these cases the genic reads were retained. Spatial tiles were designated as part of a specific sample, if >=10% of spatial barcodes of the sample match to the tile. The aligned reads were subsequently distributed across their corresponding spatial tiles per sample. Only zero Hamming distance matches between the known barcodes and the sequenced library barcodes were considered. The gene expression of tile-specific bam files was quantified with *DigitalExpression* from Drop-seq Tools (v2.5.1). The resulting quantified gene expression data was then transformed into a single h5ad file per sample, via outer merge of features (genes).

The percentage of spatially mapping reads was computed as follows. First, only uniquely mapped reads were retained, containing both genic and non-genic sequences. Quantifying gene expression via the *DigitalExpression* tool for all barcodes present in the bam file provided the total number of genic reads. Calculating gene expression in the same manner, but only for the sample barcodes, provided the total number of spatially mapping genic reads, and therefore the spatially mapping percentage.

#### Preprocessing and cell segmentation of H&E images

Tile-scan images of H&E-stained tissue sections were stitched together using the Grid/Collection stitching plugin included in Fiji 1.53t, generating a composite image of the entire section ^62^. RGB color tile-scans were converted to a HSV (Hue/Saturation/Value) image; then, the saturation channel was blurred with a Gaussian filter, and binarized with Otsu thresholding. This delivers a mask used to isolate the tissue from the background, mostly consisting of the flow cell piece. Optionally, tile-scans were further preprocessed by style transfer with a fine-tuned Contrastive Unpaired Translation (CUT) model, trained with default parameters on unpaired 512x512 pixel patches from tile-scans imaged with high noise/variance (source style), and low noise/variance (destination style) ^63^. This model was specifically applied to the metastatic lymph node H&E imaging data to equalize the style between sections and remove artifacts due to low aperture size during acquisition.

Then, nuclei were segmented with cellpose 2.2, upon fine-tuning the cyto2 model on pairs of H&E images from the metastatic lymph node sample (section 4) and manually refined segmentation masks from the same pretrained model ^26,64^. Segmentation of all images was performed with --diameter 20 (∼7 µm), --flow_threshold 2, and --cellprob_threshold –1, unless otherwise stated. Segmented nuclei were extended 10 pixels (∼3.45 µm), radially and in a non-overlapping manner, using the function `segmentation.expand_labels` from scikit-image (v0.19.3) ^65^. In samples with adipocytes (e.g., healthy human lymph node sample), a second round of segmentation with our fine-tuned cellpose model was carried out, with --diameter parameter set to 100 pixels (∼34.5 µm). Segmentation masks were similarly extended 50 pixels (∼17.25 µm) radially. To keep all small and large segmented cells, two rounds of segmentation were performed in the same image. Then, the two segmentation masks (one with and other without adipocytes) were merged by removing segmented cells from the mask with small diameter using the segmentation with large diameter as a negative binary mask. Then, the filtered small-nuclei mask and adipocyte mask were combined via AND operation.

We benchmarked the segmentation model on three randomly-selected 512x512 px regions of interest from the metastatic lymph node (section 4) tile scan. Manual nuclei segmentations, carried out with QuPath, were used as ground truth masks ^66^. Average precision at different Intersection over Union (IoU) thresholds were quantified as previously described ^67^.

#### Pairwise alignment of spatial transcriptomics data and tissue staining images

A two-step protocol was designed to align spatial transcriptomics data to tile scans of tissue staining (H&E in this study) from the same section, such that capture spot data can be aggregated into (segmented) single cells, instead of using arbitrary grids. This protocol relies on the generation of *pseudoimages* from the spatial transcriptomics data (see “Generation of Open-ST pseudoimages”), using libraries from scikit-image. First, rescaled H&E images (∼7 µm/pixel) were coarsely aligned to low-resolution (∼7 µm/pixel) *pseudoimages* of ST data via the pre-trained Detector-Free Local Feature Matching with Transformers (LoFTR) *outdoor* model, from kornia (v0.7.0). A robust transformation model was estimated with Random Sample Consensus (RANSAC). For a more precise alignment (∼1 µm error, spot resolution), fine registration was performed, leveraging feature matching on H&E images and pseudoimages with higher resolution (∼1.5 µm/pixel). Fiducial markers visible on both modalities were detected on the full-resolution images (0.345 and 0.5 µm/pixel) and appended to the features used for fine registration. The detection of fiducial marks was carried out automatically with custom YOLO-based object detection models. Likewise, the LoFTR-detected features and YOLO-detected marks were matched across modalities via brute-force and nearest neighbors matching, followed by RANSAC estimation of the transformation model. Since pairwise alignment is performed between an H&E image and ST data from the same section, rigid models were used to transform ST coordinates into H&E image space, as no distortions were expected. Finally, we used the GUI provided by our openst package (manual_pairwise_aligner_gui) to visually assess and manually refine the results of the automatic alignment pipeline. The code for alignment, GUI for visual validation, feature detection models and examples for Open-ST pairwise registration are publicly available at https://github.com/rajewsky-lab/openst.

Finally, spatial cell-by-gene expression matrices were created by aggregating the initial NxG matrix (N, capture spots; G genes) into a MxG matrix (M, segmented cells; G, genes), where the mapping of N to M takes place via the segmentation mask. That is, for all segmented regions (see “Stitching and cell segmentation of H&E-stained images”), spots falling within the spatial coordinates of a segmented region are aggregated into a single “segmented cell” with identifier equal to the cell mask label.

We assessed the accuracy of the pairwise alignment based on fiducial marks by calculating the reproducibility of two manual alignments performed on the same randomly chosen sample, (section #9 of the metastatic lymph node). Two independent annotators (L.M.S. and D.L-P.) selected at least two pairs of corresponding points per tile, using the GUI tool (manual_pairwise_aligner_gui) starting from the same automatic coarse alignment, that was performed with pairwise_aligner. Similarity matrices were computed for each tile from both lists of keypoints and applied to the coarse-aligned barcode coordinates. Lastly, the distribution of euclidean distances across annotators was calculated.

#### Generation of Open-ST pseudoimages

The generation of *pseudoimages* was tailored to distinct visualization needs, encompassing two methodologies. On the one hand, for aggregated cell-level depictions in 2D, the `pl.spatial` function from the scanpy v1.9.3 package was used.

On the other hand, higher-resolution visualizations focusing on individual transcripts (typically, at ∼100x100 µm zoom) relied on local 2D Kernel Density Estimation (KDE) of UMI-scaled spatial coordinates, for any chosen transcript. The KDE was performed with a bandwidth of ∼1 µm. For visualization, the KDE is treated as image data visualized with the microshow function from microfilm v0.2.1 package. Intensity limits were adjusted from the 5th to the 95th percentile. These renderings are useful to visualize and quantify transcript density in space, rather than the sparser raw counts.

Alternatively, raw UMI counts can be displayed as an image by producing an empty canvas defined by re-centered coordinate bounds, mapping coordinates onto the canvas, and aggregating UMIs within pixel bins. These resultant images resemble conventional pixel-based grids and can be readily subjected to standard image processing techniques.

#### 3D reconstruction of spatial transcriptomics and H&E images

The Spatial Transcriptomics ImgLib2/Imaging Project (STIM, v0.2.0) was leveraged for the alignment of the 19 Open-ST sections of the metastatic lymph node dataset. For alignment purposes, we treated sections as sequential. First, the coordinates of each pairwise-aligned and segmented h5ad file were rescaled by a factor 1:2. The coordinate and gene expression information of these datasets were converted into the n5 format, optimized for efficient image processing, via the `st-resave` function. The `st-align-pairs` function was subsequently used for pairwise alignment of three sections below and above each section (r=3), by creating image channels of the expression of prespecified genes, aggregated per cell as a Gauss rendering around centroids, parametrized with a smoothness factor. We qualitatively assessed *KRT6A*, *KRT6B*, *S100A2*, *LYZ*, *CD74*, *IGKC*, *IGHG1*, *IGHA1*, *JCHAIN*, *CD74* and *AMTN* to have high expression and spatial variation within sections. This function results in a set of feature matches between pairs of sections, filtered via affine model; `st-align-pairs` was parameterized with `--minNumInliers` 15, `--scale` 0.03, and `-sf` 4.0 (smoothness factor). Finally, we ran `st-align-global` to find affine transformations that globally minimize the distances of feature matches between sections, across the entire stack. The n5 container was then converted back to the h5ad format, for subsequent downstream analyses. We additionally transferred the resultant affine transformation matrices onto preprocessed and background-removed H&E images, rescaled to an equivalent 1:4 factor. An aligned imaging volume, used for subsequent 3D visualization, was created via the `transform.warp` function from scikit-image. This can be reproduced with the from_3d_registration program from our openst package.

#### 3D renderings

Three-dimensional rendering was used to display the two *raw* data modalities comprising Open-ST data (tissue staining images and spatial transcriptomics data), and results from downstream analyses (i.e., cell clustering).

Tissue staining renderings were generated with ParaView v5.11, from the rescaled image stack resulting from the 3D alignment (see Methods Section “3D alignment of spatial transcriptomics and H&E images”). The x-y axes were scaled 1:4 with respect to the physical dimensions, and the z axis was scaled 7:1. The x-y coordinates of the volume were clipped into a box of 4x3x0.36 mm, to remove empty areas outside of the tissue block, as well as areas with low section coverage along the z axis.

Raw or normalized gene expression of individual genes was visualized upon downscaling the expression levels per cell by a factor 1:40. Then, volumes of ∼500x500x19 voxels were generated by concatenating ∼500x500 pixel *pseudoimages* of the selected gene at each section. Upon concatenation, values were interpolated by summing the pixel-wise values from section *n-1* to section *n,* removing spatial irregularities due to uneven coverage along the z-axis. The voxelized volume data was smoothed with a gaussian filter (sigma=2), and exported as a TIFF file. Volume renders of gene expression were visualized with ParaView v5.11.

Surfaces of unsupervised, annotated clusters were generated from similarly constructed *pseudoimages* of categorical variables, with gaussian filtering and linear interpolation. This resulted in volumes of ∼500x500x19 voxels, one per annotated cluster. Within ParaView v5.11, voxel data was thresholded for isosurface extraction. Surfaces were smoothed for 1,000 iterations.

#### Clustering analysis & cell typing

We performed cell type annotation of the E13 embryonic mouse brain, human primary head and neck squamous cell carcinoma, and matched healthy and metastatic lymph node samples, by following standard practices for single-cell analysis and by using scanpy. In the case of the metastatic lymph node, the following steps were performed on one section (#4), used to build a reference annotation that was subsequently transferred to the remaining 18 sections. Data preprocessing involved applying unique molecular identifier (UMI) thresholds per segmented cell (at least 250, at most 10,000), mitochondrial count filtering (at most 10% of the counts for the mouse sample, 20% of the counts for the human sample), and retaining genes expressed in a minimum of 10 cells (Sup. Table 2). In the human primary tumor sample, we adjusted the cell filtering thresholds to at least 500, at most 10,000 UMIs per cell, and at most 15% of mitochondrial counts per cell. Then, we normalized the raw counts per segmented cell employing the scanpy functions sc.pp.normalize_total and sc.pp.log1p. Concurrently, we identified the top 2,000 highly variable genes through sc.tl.highly_variable_genes using the ’seurat’ method. Following normalization, we performed dimensionality reduction via Principal Component Analysis (PCA) on the normalized counts of the selected highly variable genes. The subsequent construction of a nearest neighbor graph in the PCA space, encompassing the first 30 principal components, was accomplished using sc.tl.neighbors. Then, we leveraged the Leiden algorithm for community clustering (sc.tl.leiden). We mapped Leiden clusters to cell types by identifying marker genes via sc.tl.rank_genes_groups, using the ‘Wilcoxon’ method, with a filter for log2FC (fold change) ≥ 1 and Bonferroni-adjusted p-value < 0.05. Subsequently, we visualized marker genes with sc.pl.rank_genes_groups_dotplot, normalizing expression per gene feature (standard_scale=‘var’). We annotated cell types from Leiden clusters by incorporating these unbiased marker genes and literature-informed markers.

For the E13 embryonic mouse head, we further clustered the cells at the forebrain, midbrain and hindbrain, separately, using the same dimensionality reduction and community clustering approach. For this, we manually generated three sets of cells based on their brain location, by filtering their x-y coordinates, and excluded cells that were annotated as chondrocytes, fibroblast, endothelial, mesenchyme, myocytes and blood. Then, clusters were annotated using literature markers. We compared our unsupervised clustering and annotation to the annotation from a reference atlas of the E13 mouse brain ^31^. We used the scVI and scANVI models from scvi-tools v1.0.2 to perform label transfer of the reference atlas to the segmented cells from our Open-ST dataset, using default settings ^68^.

For the human primary HNSCC, metastatic and healthy lymph node samples, in a secondary pipeline we refined cell type labels by aggregating subsets of cells from different samples and performing joint annotation. This allowed us to validate the robustness of our annotation regardless of the sample of origin. For the metastatic and lymph node samples, we aggregated cells not previously annotated as tumor and excluding adipocytes, which we integrated with scvi-tools v1.0.2 on raw counts of the 2,000 most variable genes. We used the default scVI model trained for 100 epochs, with 2 hidden layers and 30 latent dimensions. Similarly, we approached tumor cell sub-clustering by consolidating tumor cells from primary and metastatic samples, followed by scVI integration for batch correction on the 2,000 highly variable genes. Tumor labels were identified through Leiden cluster numbers. Lastly, cell type annotations from the metastatic lymph node section 4 were transferred across the whole dataset (rest of sections) by projecting the PCA fitted on section 4 and using a *Nearest Neighbors* classifier to map cluster labels, via sc.tl.ingest with default parameters.

#### Gene program and cell-cell communication analysis

We explored gene programs and cell-cell communication within the primary HNSCC, healthy, and metastatic lymph nodes starting from a Differential Gene Expression (DGE) Analysis, by merging the samples following a pseudobulk approach. We iteratively computed the DGE between tumor subclusters versus rest, using the sample of origin and binary indicator of cluster as design matrix, with pydeseq2 ^69^. We discerned differentially expressed genes per cell type (one versus rest) within each sample by including features with an absolute log2FC > 0.5 and an adjusted p-value < 0.05.

Subsequently, the identified differentially expressed genes per cluster underwent Gene Set Enrichment Analysis (GSEA) using the dc.get_gsea_df from the decoupler-py package. The Reactome pathway gene set signatures were employed, filtering for a FDR p-value < 0.05 and retaining programs with a NES (Normalized Enrichment Score) > 1. This analysis was executed on the various tumor cell subclusters from metastatic and primary samples, shedding light on the active pathways within unbiased clusters. We additionally validated the localization of these programs in space through AUCell scores (implemented in decoupler-py) for the respective signature gene sets.

Moving to spatial receptor-ligand analysis, we utilized the liana python package, specifying a distance of 50 µm on a manually-curated consensus database of interactions ^70^. Spatial hotspots were approached as non-negative factors, computed from normalized communication scores via the decomposition.NMF function from scikit-learn, on tumor and stromal cells. The optimal number of factors was automatically selected upon the elbow method, with the kneed package, from the reconstruction error of the inferred factors and the training data.

#### Resolution analysis

*Pseudoimages* were generated for the E13 mouse head dataset through the 2D KDE protocol at specific ROIs. The detectable scale of changes of transcript density was analyzed along manually selected lines, drawn to pass from tissue regions with *baseline* (off) to high levels (on) of expression of a specific marker. This assesses the scale at which changes from *off* to *on* can be detected, as a proxy for the effective resolution of the method (upper bound). Spatial density changes along the lines were measured via the `measure.profile_line` function from the scikit-image v0.19.3 library. Notably, for transcripts like *Ttr* (choroid plexus region), a line width of 10 pixels was employed, while for *Atoh7* (eye marker), a line width of 50 pixels was utilized. The findings of this analysis were subjected to qualitative comparison with analogous regions from a sagittal E13.5 brain, as cataloged via in-situ hybridization in the Allen Developing Mouse Brain Atlas as of 20 December 2023, for *Neurod6* (Image 2), Pbx3 (Image 2), *Tubb3* (image 7), *Atoh7* (image 1), *Ttr* (image 7), *Cnr1* (Image 3), *Dbi* (Images 1 and 4), *Eomes* (Image 4), *Tbr1* (Image 5), *Pax6* (Image 2), *Shox2* (Image 8), *Npy* (Image 8), *Six3* (Image 10), *Enc1* (Images 3 and 5), *Nes* (Image 6), *Htr2c* (Image 5), *Cyp26b1* (Image 6), and *Foxp2* (Image 5).

#### Subcellular localization analysis

To study the average subcellular localization patterns of transcripts, the adult mouse hippocampus dataset, consisting of transcript locations at ∼0.6 µm spots and the pairwise-aligned segmentation mask, was partitioned into a regular grid of 70x70 µm squares. The cellular density of all patches was assessed by computing the proportion of nuclear segmentation mask coverage over total patch area. Patches were further categorized into three tiers: 10-15%, 15-30%, and 30-50% mask coverage, or nuclei density.

The same process was applied within each density group, for three transcript or transcript families: *Malat1* (nuclear), *mt-\** (all mitochondrially-encoded transcripts), and *mt-Tt* (a transcript with similar total UMIs to *Malat1*). A 3.5 µm extension surrounding the nuclear segmentation mask was enacted, and for distinct cells, distance transforms were calculated utilizing the ndimage.distance_transform_edt function from scipy v1.10.0 ^71^. Through a linear rescaling and inversion, these distance transform values were mapped to the continuous [0, 2] interval, wherein 0-1 corresponds to the nuclear core and 1-2 pertains to the extended boundary. This allows for a standardized comparison across cells. For both *Malat1* and mitochondrial transcripts, the coordinates were projected onto the rescaled distance transform, producing the “observed” distributions that encapsulate the spatial behavior of the transcripts within cells.

An "uniform" distribution was generated by shuffling the location of transcripts prior to projection to the distance transform, for later statistical comparisons. Kernel density estimation was performed, utilizing 40 bins within the [0, 2] interval, to approximate a smooth distribution shape. Measurements from all cells were aggregated, and an observed-to-uniform ratio was computed for each transcript or transcript family. These ratios were visualized within standardized cells, via polar projection of the density estimates at each rescaled distance value, equivalent to radii of a standardized (circular) cell. As an additional validation, the histogram and kernel density estimate of *Malat1* and *mt-\** counts respect to the distance to nuclear edges was computed. To this end, we projected the points into a distance-transformed mask, where zero values correspond to the nuclear edge (pixels delimiting the boundary between nuclear segmentation and background), negative values point towards the centroid of the nucleus, and positive values diverge from the edge to the outside of nuclei.

#### Crosstalk and spatial bias analysis

We examined the biases in transcriptomic profiles of cells by the spatial biases in transcript capture by defining crosstalk as how the expression from a cell of population *a* has components of cell of population *b*, depending on their pairwise spatial distance. We define a crosstalk coefficient 𝑘_*ab*_ as the expression of markers from cell type *b* measured at cells *a*:

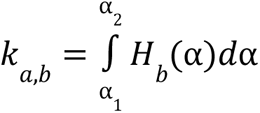

Where α_1_ and α_2_ are the expression (or gene set score) limits for population *a*, and 𝐻*_b_* is a function that returns the amount of expression (or gene set score) for population *b*, measured at level α in population *a*.

We selected the E13 embryonic mouse head sample, and we subsetted the data to the cells of the clusters labeled as ’Blood’ and ’Fibroblast’ due to their widespread presence across the sample and their distinct marker expression, determined through unbiased clustering (detailed in the ’Clustering’ section). We performed the following analysis on specific cell pairs using either a segmentation mask or a hexagonal grid with a 7 µm radius: first, we calculated marker gene set scores for these clusters using the scanpy function `scanpy.tl.score_genes`; then, we calculated pairwise spatial distances between ’Blood’ and ’Fibroblast’ cells. Employing a spatial filtering criterion of a center-to-center distance under 20 µm, we generated 2D contour plots of kernel density estimates, portraying gene set scores of ’Fibroblast’ and ’Blood’ markers in the x and y axes grouping the density clouds based on the label of the cells of origin. In practice, the crosstalk coefficient 𝑘_𝑎𝑏_ measures the overlap of the two KDE clouds. Finally, to extend the analysis, we performed this analysis on a dataset with simulated lateral diffusion for individual transcript coordinates, with an equivalent diffusion coefficient D= 1 µm/s^2^ for 100 seconds, via random walk (dt = 1 s).

#### Neighborhood analysis

We quantified the spatial relationships between distinct cell types in primary HNSCC and metastatic samples by studying local cellular neighborhoods. Utilizing a Delaunay triangulation, we computed interaction graphs where two cells are connected if they share an edge. This provided a quantitative measure of the frequency of cell type neighborhoods, that further allows to assess the enrichment of neighborhoods via permutation tests. For this purpose, we employed the squidpy and CellCharter frameworks ^72,73^ and binarized the asymmetric neighborhood enrichment matrix to keep values with p-value < 0.05 and relative enrichment > 0, i.e., we ignore neighborhood *depletion* events.

Tumor boundary analysis

To define the boundary between the tumor mass and lymph node in the metastatic sample, we start from the radius neighborhood graph and define edges for pairs of cells with center-to-center distance smaller than 50 µm. The value for a node is then computed as the number of edges connecting to a cell type from the lymph node, if the node is a tumor cell, and vice versa. The set of these values along all nodes is the boundary strength 𝑆. To calculate the correlation of two signals in space, e.g. gene expression to a boundary, we compute a value (𝐼) inspired by the bivariate counterpart of Moran’s I:

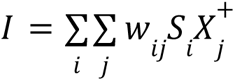

where 𝑤 is the weighted adjacency matrix from the spatial graph, *X*^+^ is the query signal, and 𝑆 is the boundary strength. *X*^+^ results from setting negative values from 𝑋, the normalized and z-scored gene expression matrix, to zeroes. We analyzed the 9,000 genes with highest mean expression, and reported the ranking of top-50 genes with highest *I*\* values, computed as *I*\*= 1^+^(𝐼)/1^−^(𝐼), where 1^𝑐^ is an indicator function setting ones where the values fulfill the comparison term 𝑐. We computed the score of the set of top-50 genes sorted by highest 𝐼* values with `scanpy.tl.score_genes` and visualized it in space to validate the enrichments.

#### Flow cell imperfection analysis

To quantify macroscopic imperfections exceeding 2 µm, an iterative approach was employed, separately for each tile at the the top and bottom parts of the S4 flow cell. First, a *pseudoimage* of the flow cell was generated by rescaling the coordinates of individual capture areas sourced from spatial barcodes (see Methods Section “*Quality control and spatial reconstruction of barcode sequencing*”) by a factor of 0.1. This rescaling aligns consecutive spatial points to an interval of one unit length (1, 2, 3… corresponding to sequential positions), resulting in a 4430x4600 pixel image. Gaussian blurring, with a sigma of 5, was applied to the *pseudoimage*, followed by Otsu thresholding. The outcome is a segmentation of space filled by barcoded spots, wherein pixel values are either 1 (presence of spatial barcodes) or 0 (absence). By implementing this procedure, high-frequency fluctuations were suppressed, isolating larger-scale spatial voids.

This sequence of operations was replicated for all tiles within the flow cell. Subsequently, the proportion of pixels with a value of 0 within the segmentation mask is computed against the total pixel count. This value, termed the ’percentage of irregularities,’ characterizes regions with deviations in the flow cell surface. Such irregularities may stem from diverse factors, including fiducial marks whose contrast is enhanced following blur and thresholding.

#### Benchmarking analysis

Slide-seq and Seq-Scope data were downloaded from the Gene Expression Omnibus (accession codes: GSE197353 and GSE169706, samples: GSM5915059 and GSM5212844, respectively), Visium data from the [10x Genomics website](https://www.10xgenomics.com/resources/datasets/mouse-brain-serial-section-2-sagittal-anterior-1-standard) and Stereo-seq data from the CNGB Nucleotide Sequence Archive (experiment ID: CNX0422301) ^10,16,74^. All barcode files were formatted to: bc \t x_pos \t y_pos. To reduce computational costs for the Stereo-seq data the barcode file was split into chunks of 10 million barcodes each and 1.5 billion of the 5.2 billion total reads were randomly sampled. All datasets were processed using Spacemake (v0.7.3) with the parameters as listed in Supplementary Note 2 ^27^. Seq-Scope, Stereo-seq and Open-ST data were binned into hexagons with 7 μm edge length, resulting in an area of 127.3 μm^2^ per spatial unit. To remove background signal and low quality beads cutoffs of 20, 180, 5000, 200, 700, 700, 350, 600 UMIs per spatial unit were applied to Slide-seq, Seq-Scope, Visium, Stereo-seq, and Open-ST mouse brain, healthy lymph node, metastatic lymph node, primary tumor data, respectively. A field of view with representative quality was chosen to focus on the desired tissue area. Assuming sizes of 78.5 μm^2^, 2,376 μm^2^ and 127.3 μm^2^ for Slide-seq, Visium, and Seq-Scope, Stereo-seq, Open-ST, respectively, spatial units within the field of view were randomly sampled such that the total covered area corresponds to 1mm^2^, resulting in 12,738, 420 and 7,855 spatial units, respectively. For each dataset, reads belonging to the sampled spatial units were extracted from the output .bam file created by spacemake with samtools (v1.17) view -D and afterwards shuffled using samtools collate. The shuffled reads were downsampled in steps of 4 million reads up to a maximum of 40 million. All the downsampling files were processed using the DigitalExpression function from Drop-seq tools (v2.5.1) to obtain the genic reads and transcript numbers per barcode from the resulting summary files. These were summed to get the total number of genic reads and transcripts in the 1 mm^2^ area. The total number of transcripts was divided by 10,000 to get the average number on a 100 μm^2^ area. The total number of reads was computed as: total reads = total genic reads / (1 - PhiX mapping%) / (1 - rRNA mapping%) / spatial mapping%, where PhiX, rRNA and spatial mapping% where obtained as specified in *Quantification*.

## Additional resources

We have set up an online resource with detailed descriptions of all experimental and computational steps: https://rajewsky-lab.github.io/openst.

## Key resources table

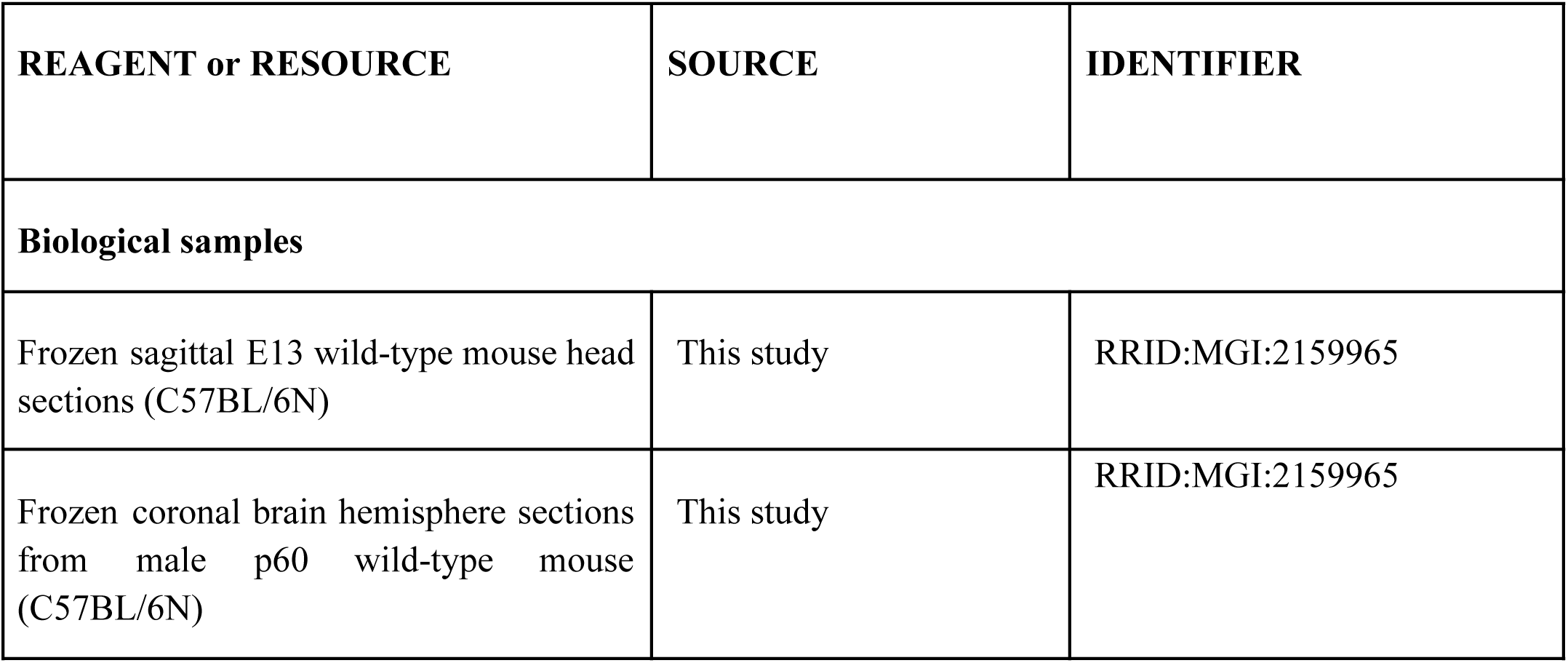
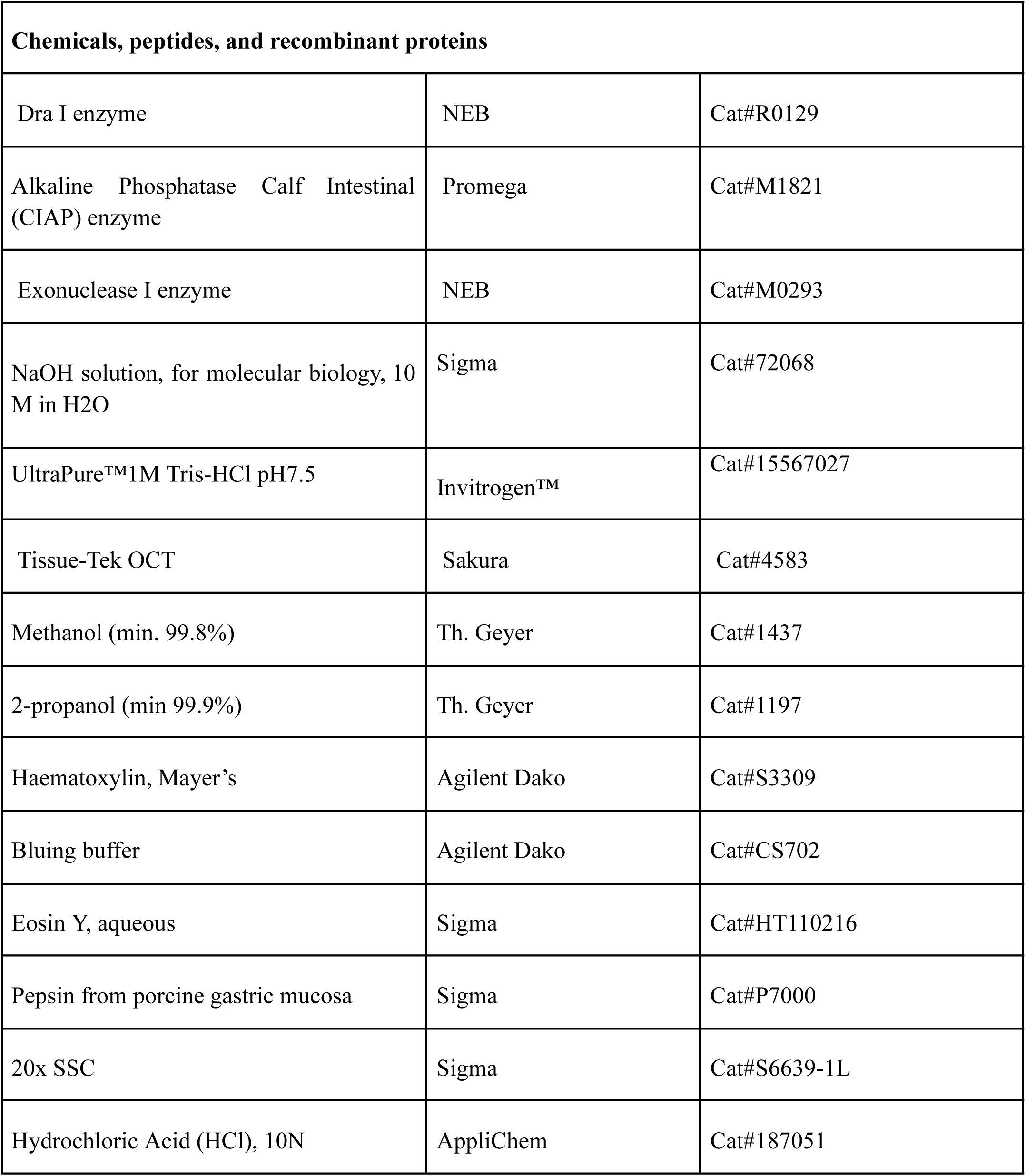
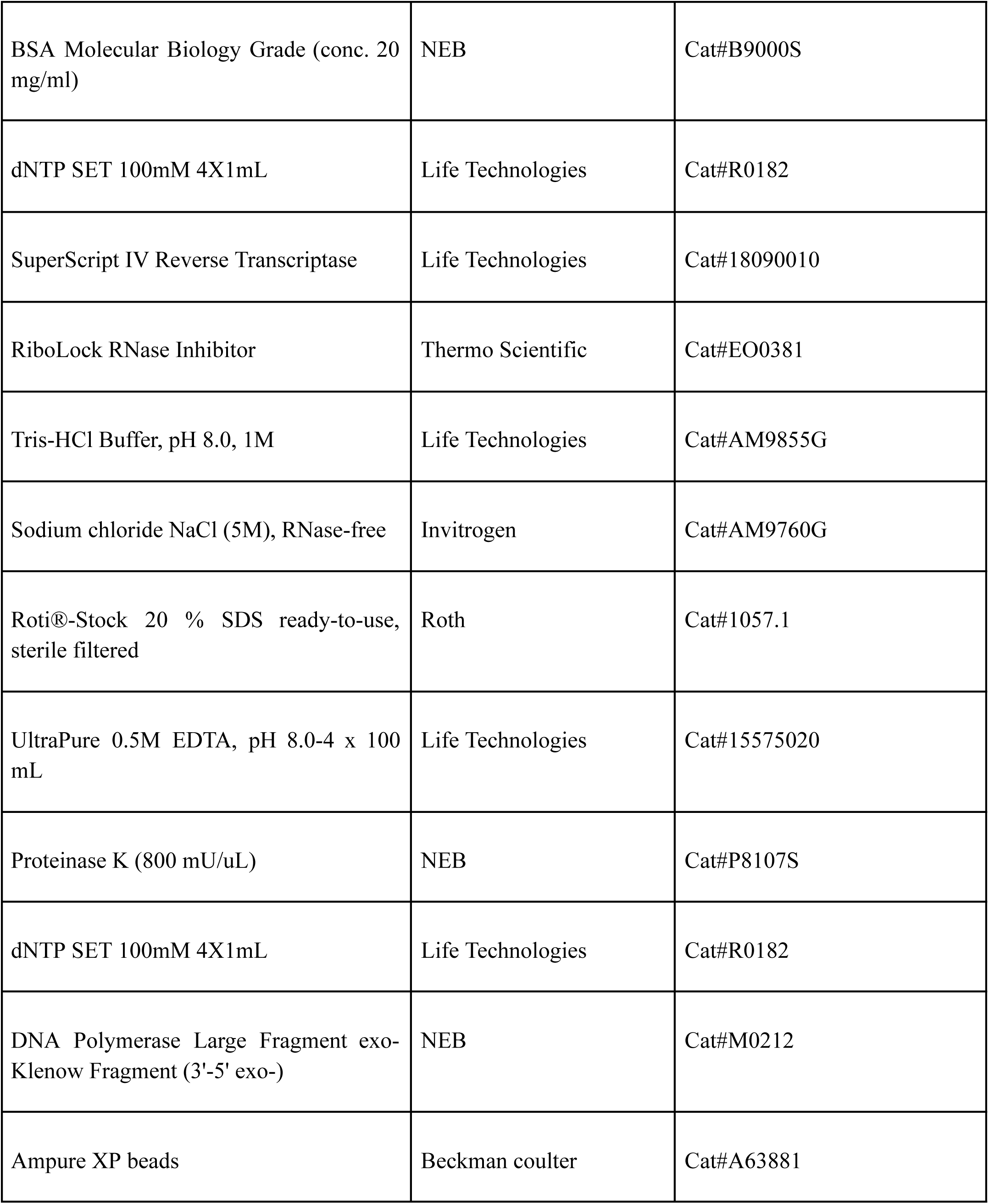
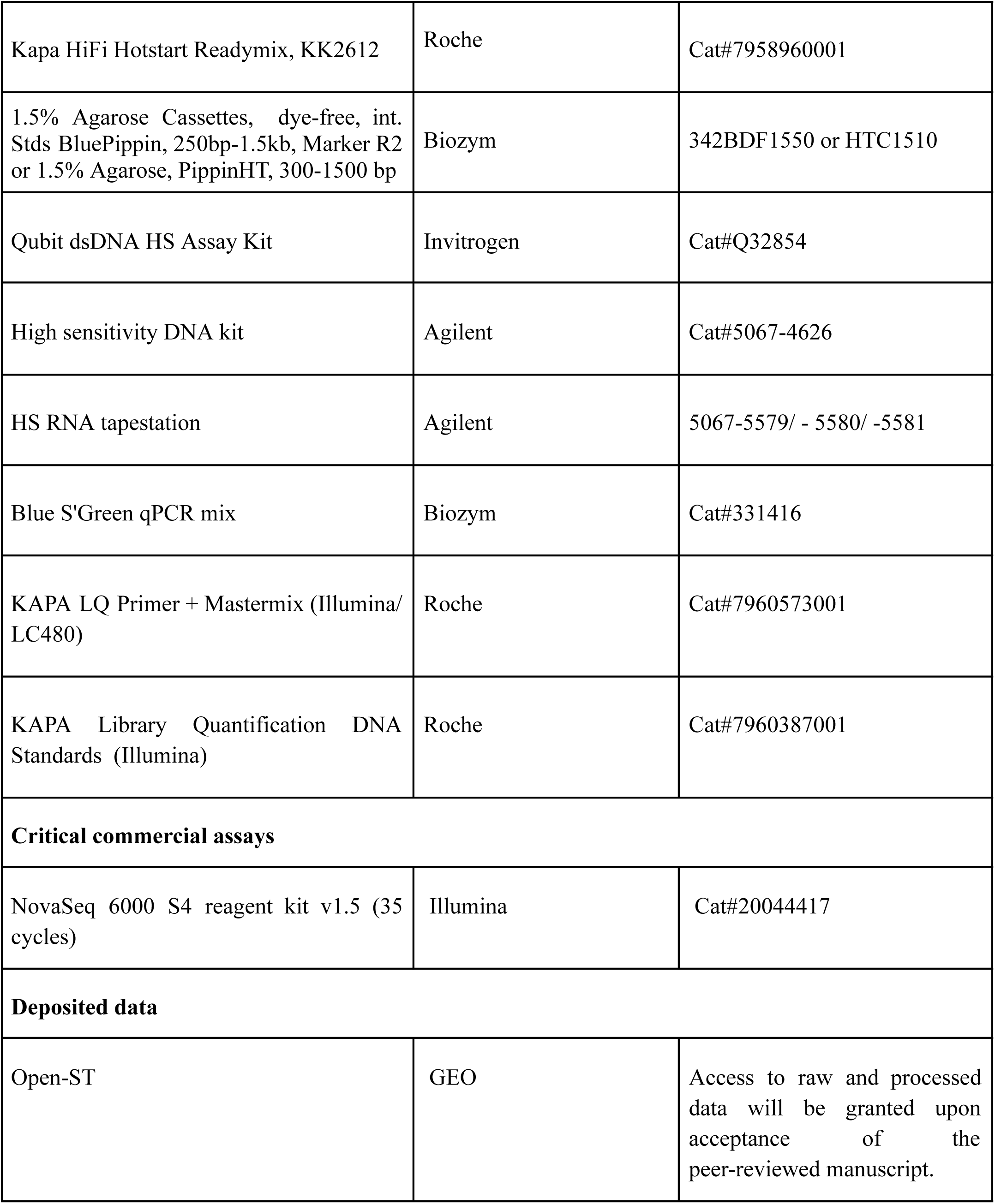
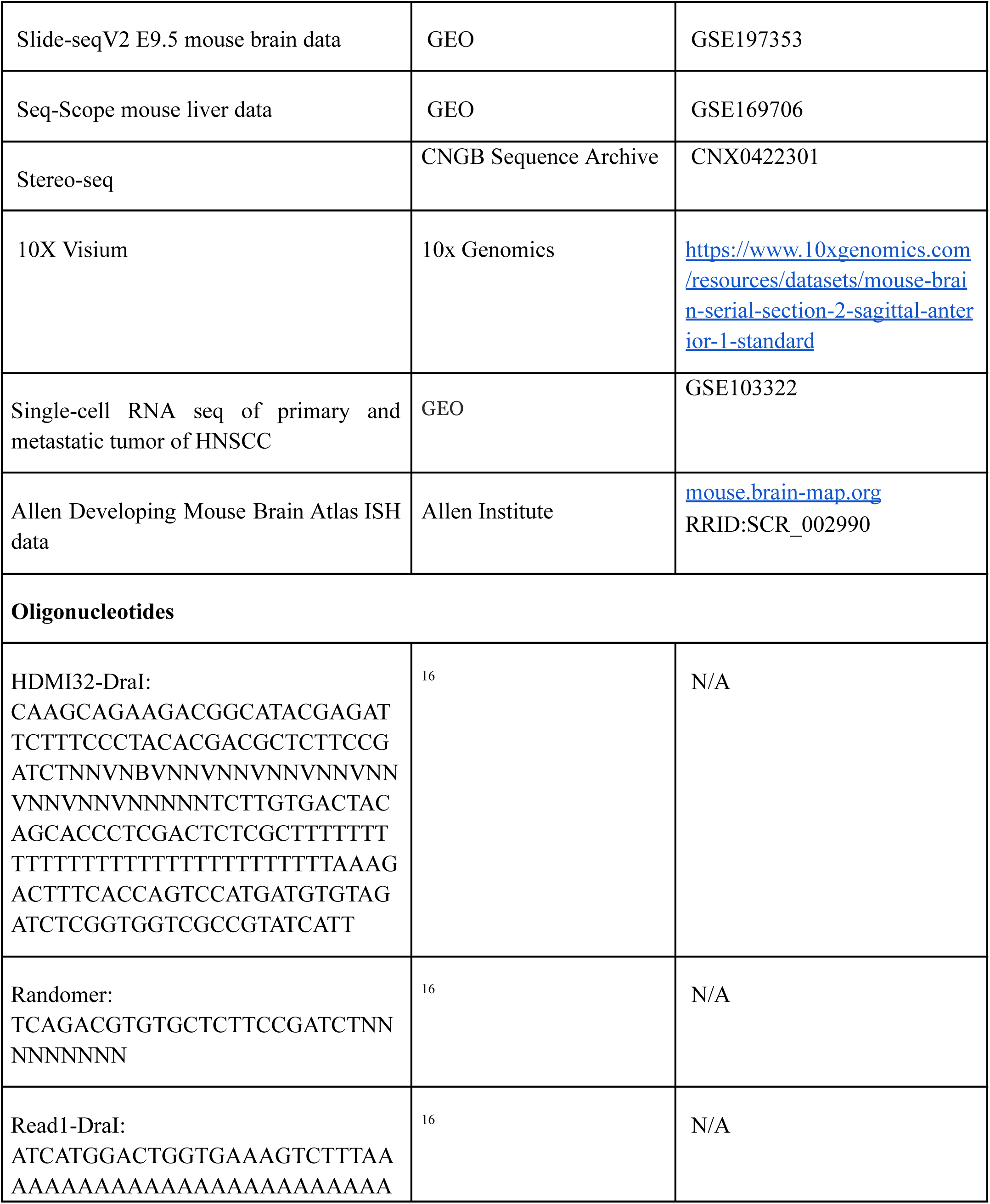
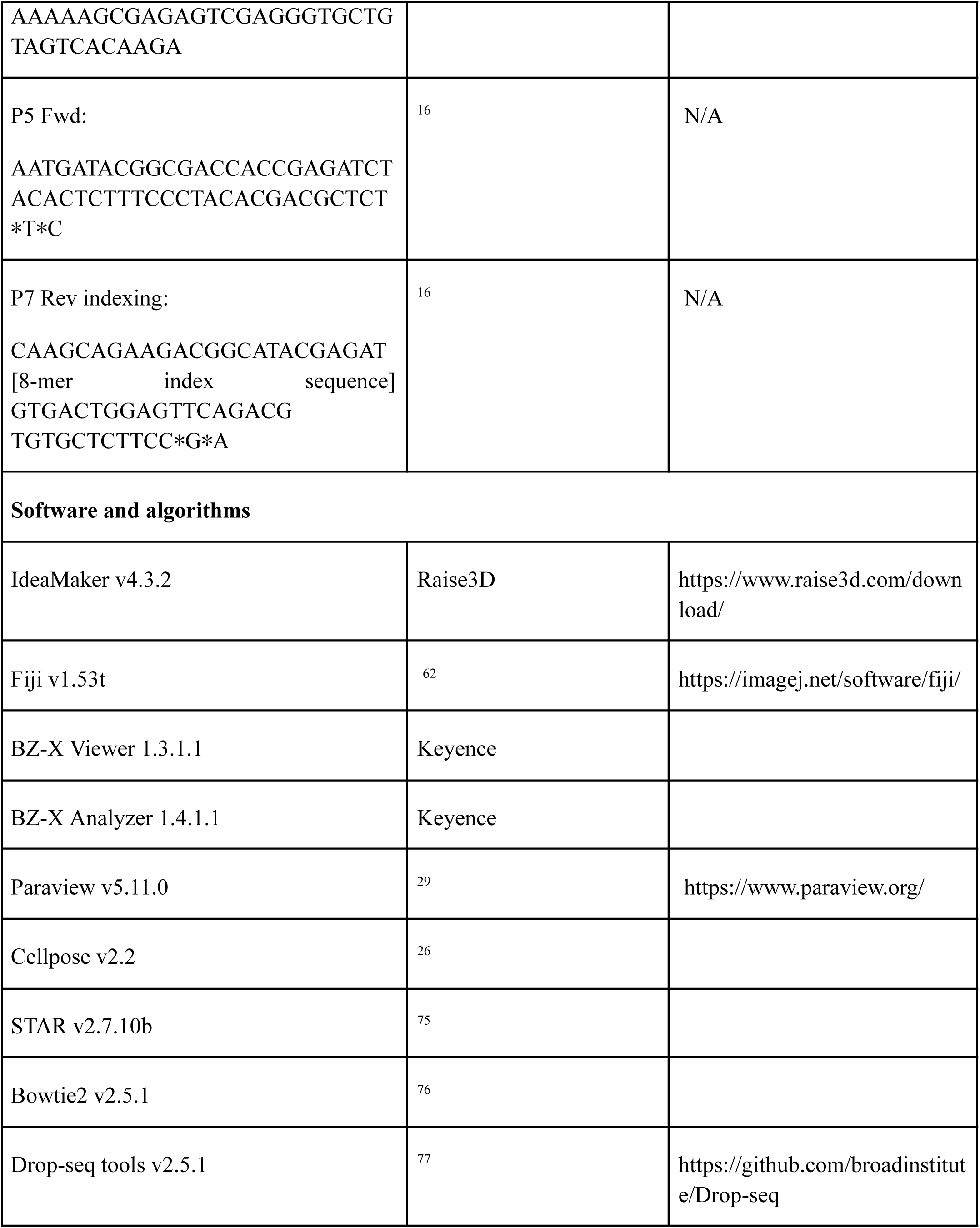
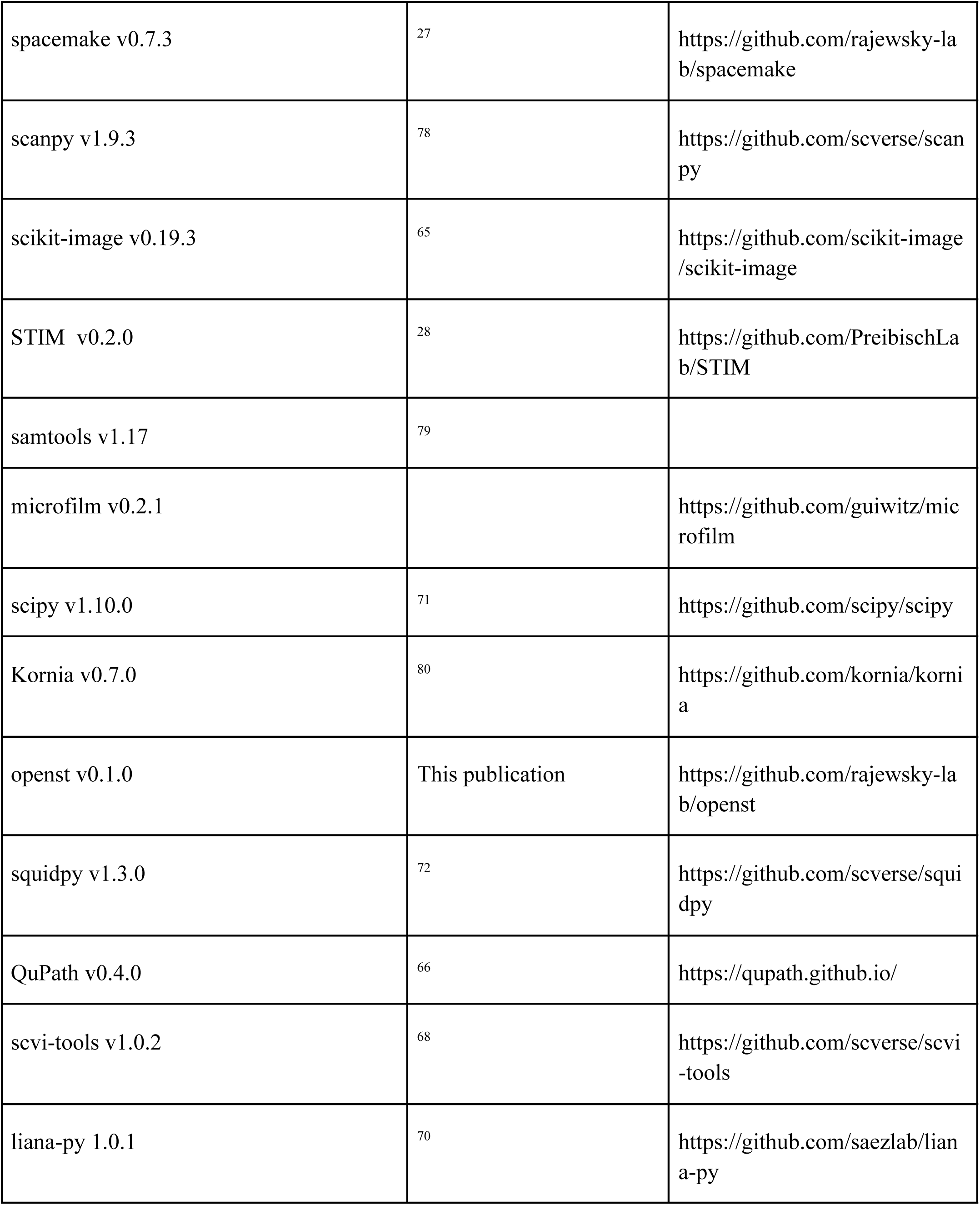
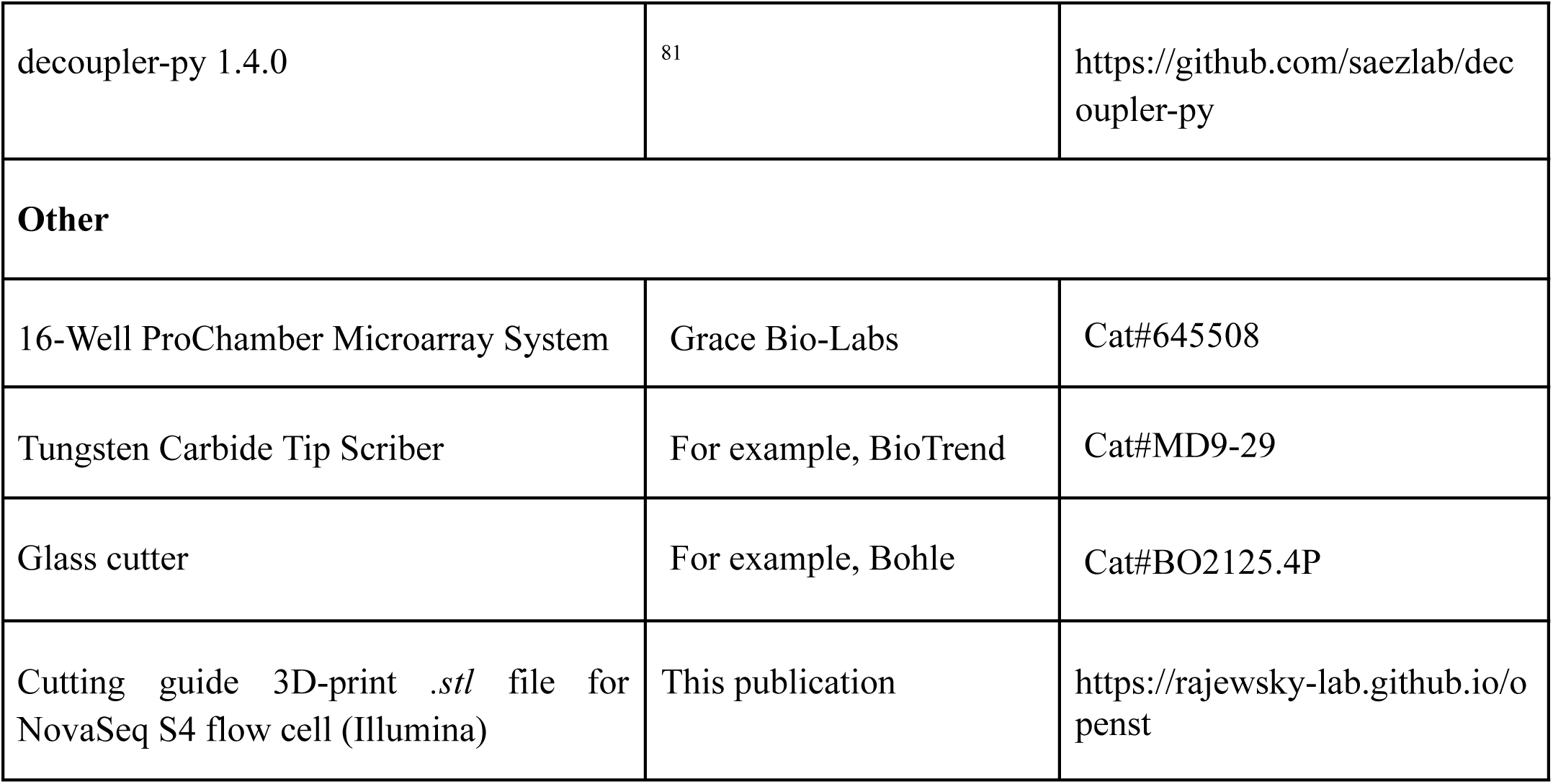

---

**Figure S1.**
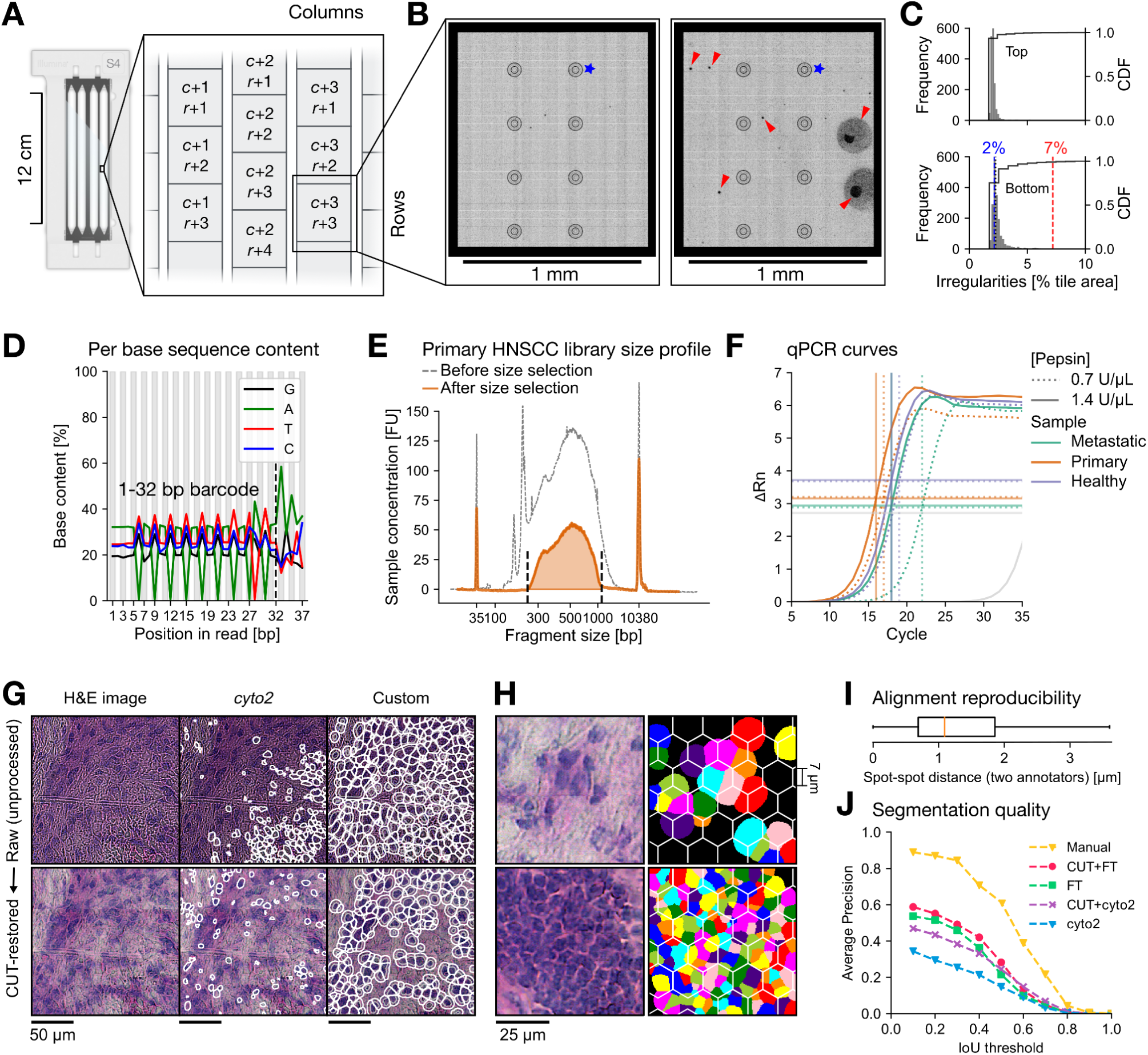
Open-ST: quality control and image processing, related to Figure 1. **(A)** A NovaSeq S4 flow cell consists of 4 lanes with a top and bottom surface, each with 6 columns (c) and 78 rows (r) per lane, totalling 3,744 tiles. Tiles are discrete sections of the flow cell imaged during sequencing. Distance between tiles is 55.5 µm in the x- and 5.3 µm in the y-axis. **(B)** Representative example of tiles with average number of irregularities (left) and higher number of irregularities (right). Blue star: space without spots due to fiducial markers; red arrowhead: imperfections that may result from flow cell manufacturing errors (small black dots), bubbles during sequencing (medium black areas), or dust obstructing the imaging (large black areas). **(C)** Frequency of irregularities (>2 µm) on the flow cell fc_1 from the 1st sequencing. The top flow cell surface contains fewer irregularities (CDF, cumulative density function). **(D)** Per base sequence content of 1st sequencing of fc_1. Bases 1 to 32 correspond to the barcodes, with drops in “A” or “T” at expected sites in sequence (B or V in IUPAC nucleotide code). **(E)** Automated electrophoresis profiles of the primary HNSCC library before and after size-selection. Fluorescent units (FU) relate to bioanalyzer input and not total sample concentration. Peaks at 35 and 10,380 bp are the upper and lower markers. **(F)** Comparison of tissue permeabilization conditions via qPCR assay (Methods). Example data with two pepsin concentrations tested per tissue type: permeabilization with higher concentration captured more mRNA. **(G)** Image restoration and segmentation pipeline: raw images after stitching (upper), and restored images after applying a Contrastive Unpaired Translation (CUT) model (Methods). Left: H&E images used as input; middle: segmentation masks generated by Cellpose 2.0; right: segmentation masks via our fine-tuned version (Methods). **(H)** Robustness of segmentation for different cell densities; two regions with ∼65% (upper) and ∼99% density (bottom) shown for the metastatic lymph node. CUT-restored images (left) were segmented with our fine-tuned (FT) model (right). Each segmented cell (nucleus with a 3.5 µm extension) is depicted with a different color. A regular lattice of hexagons (7 µm side) is overlaid onto the segmentation masks, illustrating the difference between data meshing and segmentation. **(I)** Accuracy of the spatial, pairwise alignment between staining image and barcoded spots, measured as the pairwise euclidean distance of barcoded spot coordinates after two independent manual selections of correspondences (Methods). **(J)** Benchmark of the segmentation models, measured as the average precision of the restoration+segmentation models at different intersection over union (IoU) thresholds (Methods).

**Figure S2.**
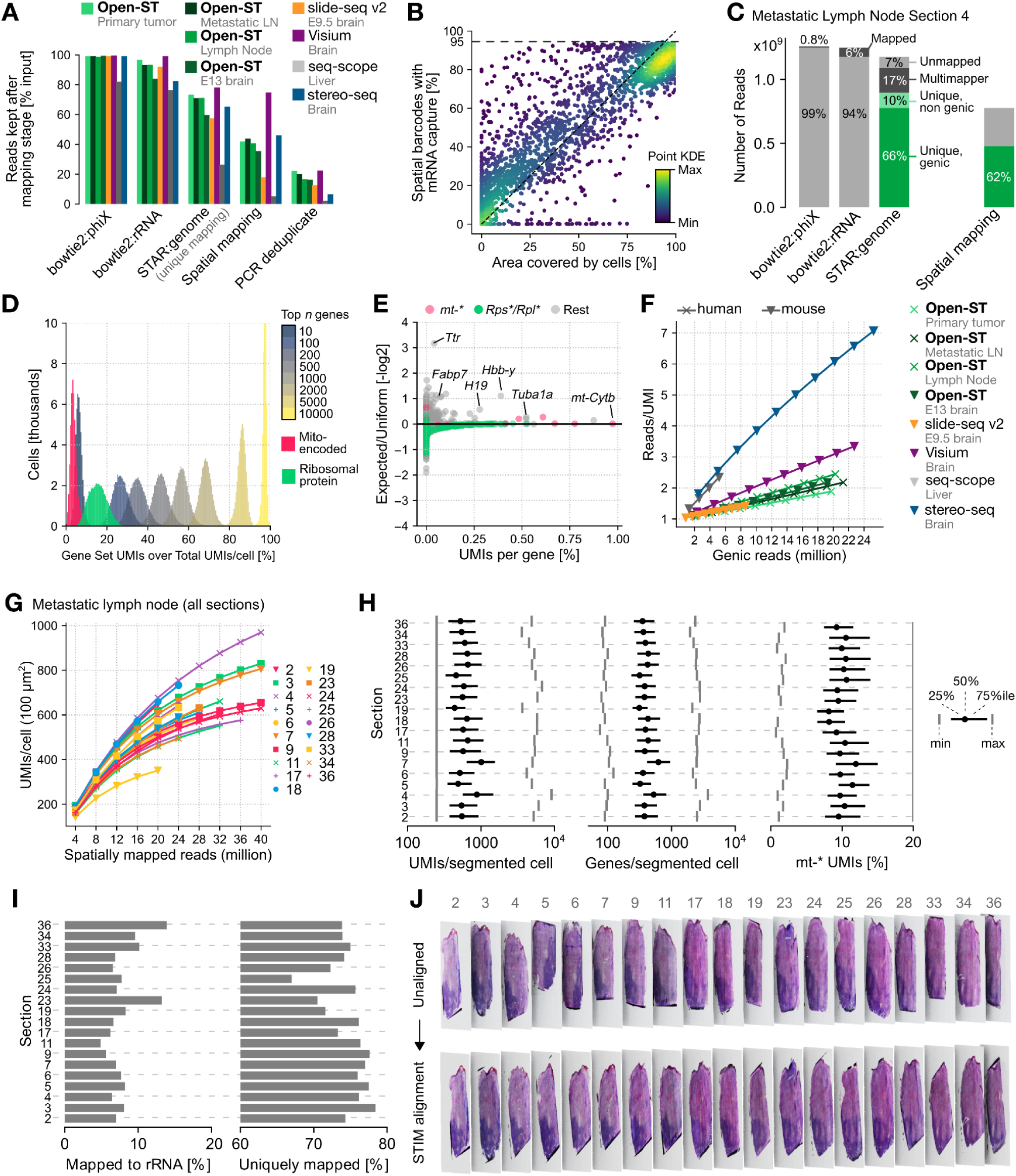
Reproducibility and performance of Open-ST, related to Figure 2. **(A)** Reads retained across different technology datasets after discarding those aligning to PhiX and rRNA, multi-mappers, reads not associated with a spatial barcode, and removing PCR duplicates. Coloring as in (F). **(B)** Percentage of spatial barcodes with captured transcripts relative to the area covered by cells at non-overlapping ∼5,000 µm^2^ square regions. Diagonal dashed line: y=x; horizontal dashed line: y_max_ across the dataset. Color gradient shows point density. **(C)** Stacked barplot of input reads filtered out during two-stage alignment against PhiX and human rRNA, summarized STAR mapping statistics from the remaining reads, and percentage of uniquely mapping genic reads with matching spatial barcodes, for the metastatic lymph node section 4 dataset. **(D)** Percentage of total unique molecular identifiers (UMI) per segmented cell accounting for several gene sets: top *n* genes by total UMIs, mitochondrially-encoded transcripts (*mt*-*), and ribosomal protein transcripts (*Rps*\*/*Rpl*\*). **(E)** Transcriptomic information measured as the expected/uniform ratio of cells expressing a gene, over the total UMIs of that gene across cells. **(F)** Library complexity measured as reads/UMIs ratio over number of genic reads across samples. **(G)** Sequencing investment measured as UMIs/cell per 100μm^2^ over spatially mapped reads for all 19 sections of the metastatic lymph node sample. **(H)** Distribution of UMIs and genes captured per segmented cell, as well as % UMIs mapping to the mitochondrial genome, across all 19 sections of the metastatic lymph node. **(I)** Left: % reads mapping to rRNA after excluding PhiX-mapping reads; right: % reads uniquely mapping to the genome after excluding rRNA-mapping reads. Data shown across all 19 sections of the metastatic lymph node. **(J)** Stacked 3D rendering of H&E images before (top) and after global alignment with STIM (bottom).

**Figure S3.**
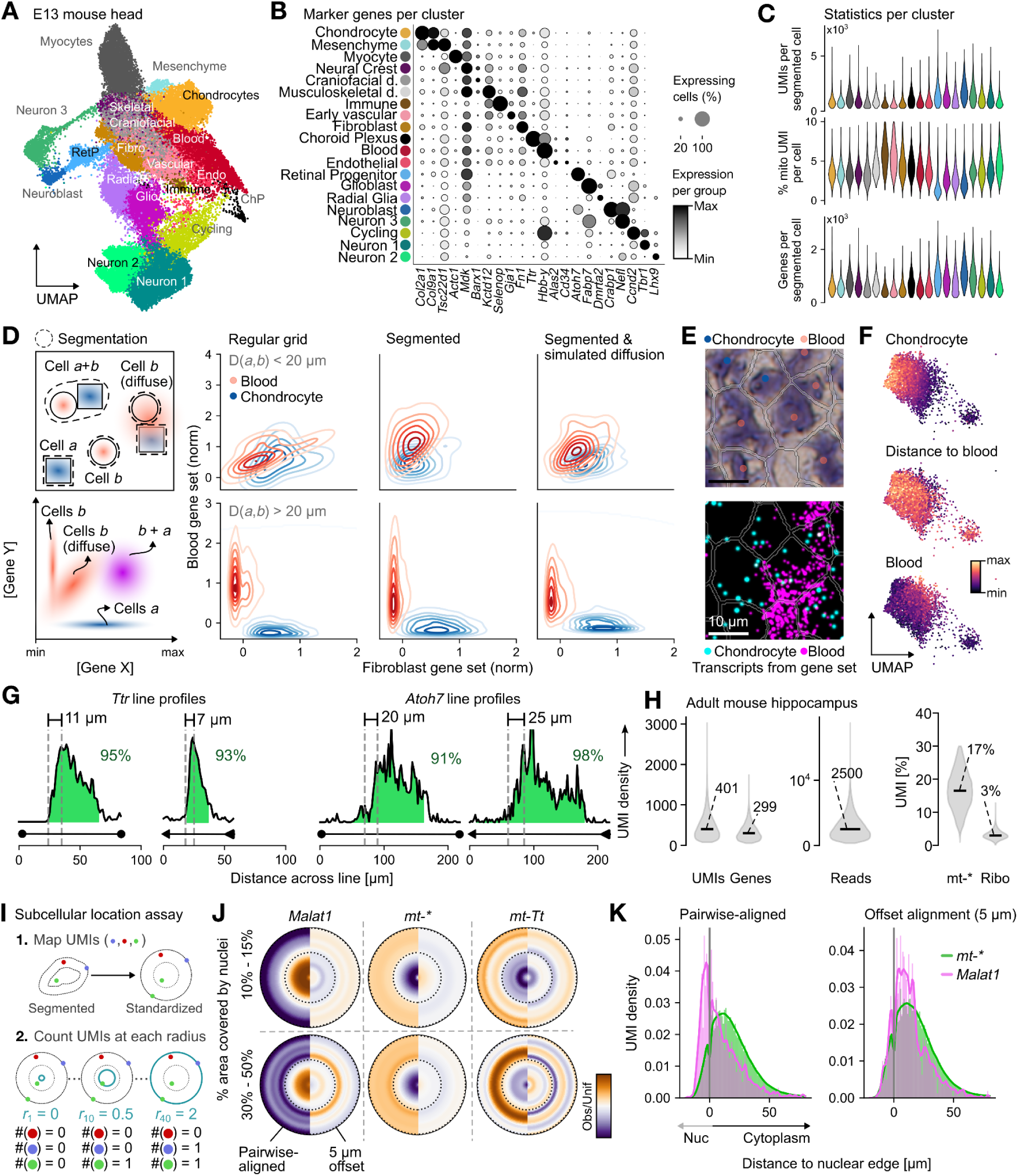
Localized transcript capture yields accurate annotation of mouse head cell types, related to Figure 3. **(A)** UMAP visualization of the E13 mouse head clusters. **(B)** Normalized expression of selected marker genes across the annotated Leiden clusters of the mouse E13 head. **(C)** Distributions of UMIs, % of mitochondria-encoded transcripts and genes per segmented cell for the mouse E13 head. **(D)** Crosstalk analysis between ‘Blood’ and ‘Fibroblast’ clusters. Left, top: sketch of two cell types (*a*: circle, *b*: square) with their expression of marker genes in space (gene Y: orange, gene X: blue), alongside dashed outlines indicating cell segmentation. Left, bottom: measurement of gene expression of X and Y, in cells of type *a* and *b*. Right: quantifications of the ‘Blood’ and ‘Fibroblast’ marker gene sets at cells. From left to right, cells are delimited via a regular grid (hexagons of 7 µm side), or via nuclear segmentation masks with radial extension. Additionally, lateral diffusion of transcripts is simulated and grouped into the same segmentation masks. Measurements focus on cells with neighboring ‘Blood’ or ‘Fibroblast’ cells at less (top) or more (bottom) than 20 µm center-to-center distance (Methods). **(E)** A region of the E13 mouse head with ‘Blood’ and ‘Chondrocyte’ cells in close proximity on the H&E staining with the segmentation mask outline (top), and the expression of marker gene sets in space (bottom). **(F)** UMAP for the subset of ‘Fibroblast’ cells. From top to bottom, gene set scores for fibroblast markers, distance of cells to the nearest cell classified as ‘Blood’ in space, and gene set score of ‘Blood’ markers. **(G)** Linear intensity profile of selected regions shown in Figure 3F, with vertical lines indicating local maxima and minima used for distance measurement. Green: Proportion of area under the curve within tissue boundaries (green) is given (Methods). **(H)** Distributions of UMI, gene and read counts per segmented cell in a coronal section of the adult mouse hippocampus hemisphere (left) and the relative distribution of UMIs corresponding to mitochondria-encoded or ribosomal proteins (right). **(I)** Outline of the method for mapping and counting spatial UMIs at a standardized (circular) cell. **(J)** Observed over uniform UMI per radius (*n_radii_* = 40) at two cellular densities, for *Malat1,* all mitochondrially encoded transcripts and *mt-Tt* (Methods). **(K)** Distribution of UMI counts with respect to the nuclear edge. Left: spatial distribution of UMIs for *Malat1* and mitochondria-encoded transcripts. Right: distribution profile after applying a two-dimensional offset of 5 μm to the pairwise-aligned spatial coordinates. Negative distances correspond to space within the nucleus (Methods).

**Figure S4.**
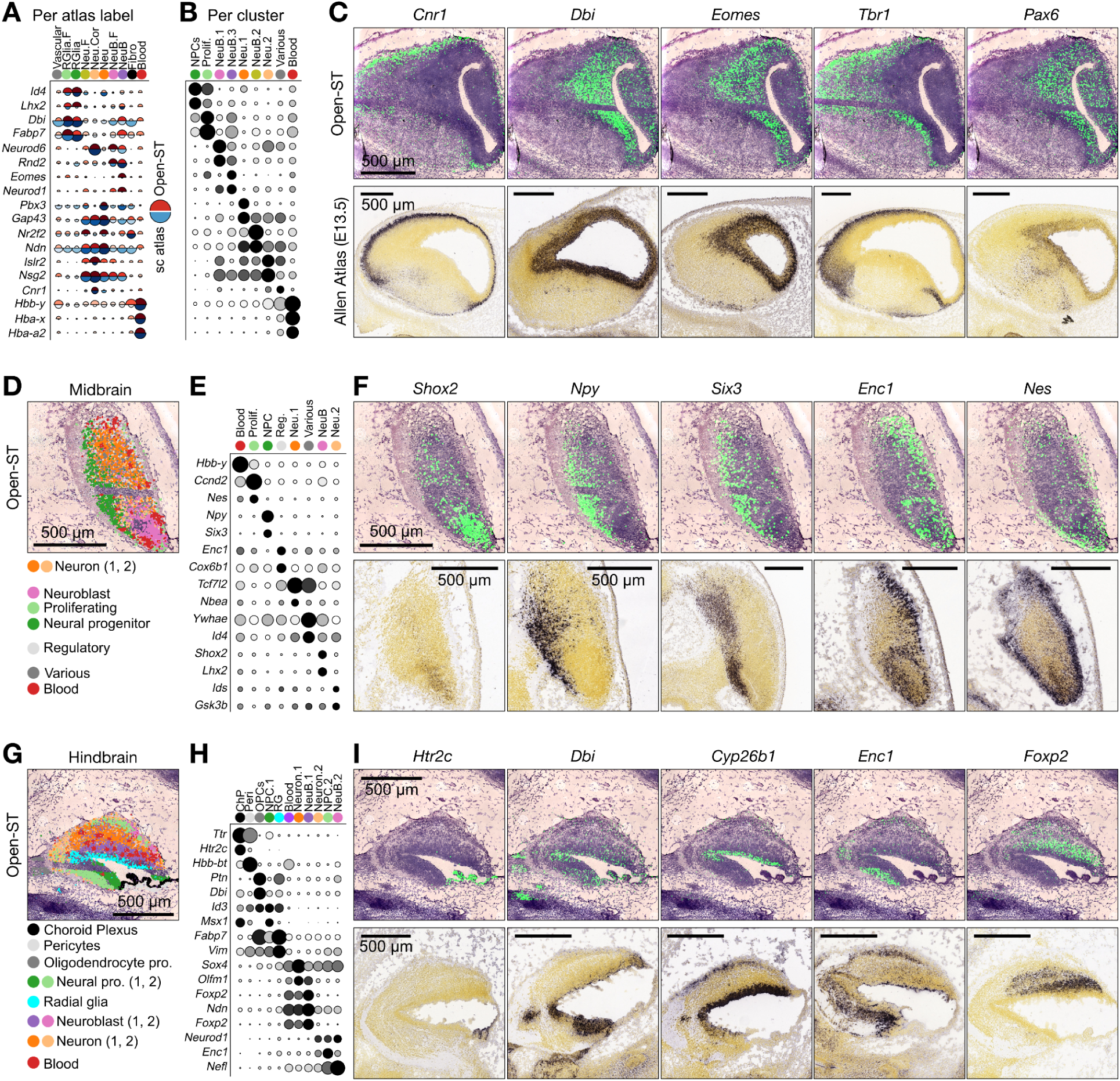
Marker gene localization in E13 murine brain regions, related to Figure 3. **(A)** Normalized expression of selected marker genes of the E13 mouse forebrain in the segmented cells of the Open-ST data (red, top) and in the cells from the E13 reference atlas (blue, bottom), grouped by the labels in the published atlas (Methods)^31^. **(B)** Normalized expression of selected marker genes of the E13 mouse forebrain subclusters, grouped by the cluster annotation labels. Genes shown in (C) are indicated in bold. **(C)** Localized capture of selected marker genes in the E13 mouse forebrain profiled with Open-ST (top) compared to *in situ* hybridization images of the E13.5 mouse from the Allen Developing Mouse Brain Atlas (bottom). High expression is colored in green (black) for Open-ST (Allen Atlas). **(D)** Spatial distribution of E13 mouse midbrain subclusters (top) and corresponding region with annotation of morphological regions from the Allen Developing Mouse Brain Atlas (bottom). **(E-F)** As in (B-C), but for the midbrain. **(G-I)** As in (D-F), but for the hindbrain.

**Supplementary figure 5.**
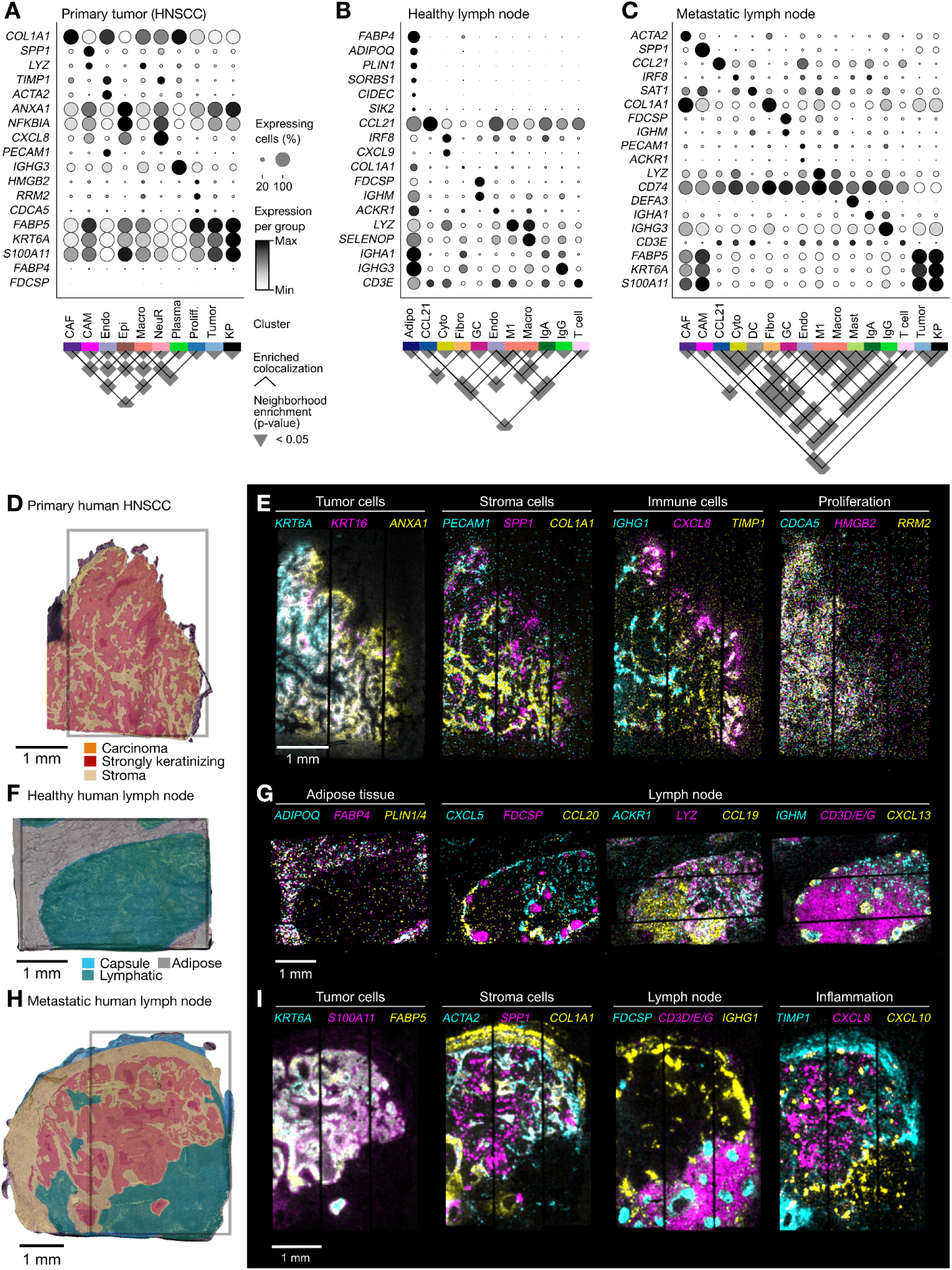
Transcriptomic clusters are consistent with pathologist’s annotation, related to Figure 4. **(A-C)** Normalized expression of selected marker genes (top) and neighborhood enrichment (bottom) across the annotated Leiden clusters of the primary HNSCC (A), the healthy lymph node (B) and the metastatic lymph node (C). **(D)** Pathologist’s manual annotation of tissue domains superimposed on H&E image of the HNSCC section. The rectangle indicates tissue area processed with Open-ST. **(E)** Spatial distribution of gene expression in the primary HNSCC section. Representative gene markers depict the molecular landscape of distinct tissue regions. Visualized as a merged representation of intensity channels, the coexpression of genes manifests as a ’sum’ of different colors (white) (Methods). **(F)** As in (D), but in the healthy lymph node. **(G)** As in (E), but in the healthy lymph node. **(H)** As in (D), but in the metastatic lymph node section 4. **(I)** As in (E), but in the metastatic lymph node section 4. Prolif: proliferating; KP: keratin pearl; NeuR: neutrophil-recruiting; Epi: epithelial; CAM: cancer-associated macrophage; Macro: macrophage; Endo: endothelial; CAF: cancer-associated fibroblast; GC: germinal center; Fibro: fibroblast; Cyto: cytotoxic T-cell; M1: M1 macrophage; Fibro: fibroblast; Endo: endothelial; Adipo: adipocyte

**Figure S6.**
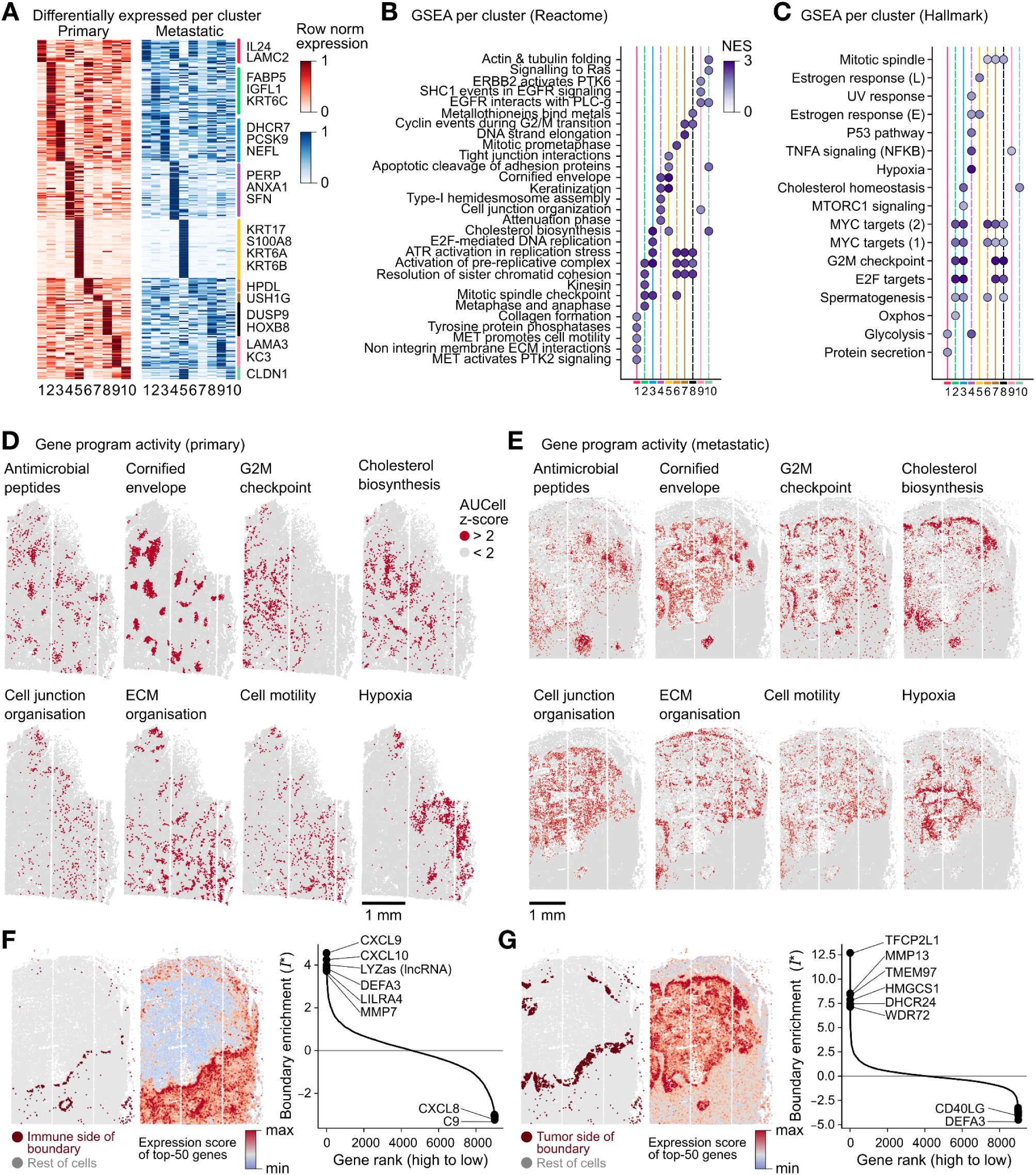
Tumor subclusters are linked to distinct spatial activity of gene programs. **(A)** Normalized expression of selected marker genes across the annotated Leiden clusters of tumor cells in the primary HNSCC (red) and the metastatic lymph node (blue). **(B)** Top five reactome pathways sorted by Normalized Enrichment Score per subclusters of the primary and metastatic HNSCC tumor cells, with NES > 1 and FDR-adjusted p-value < 0.05 (Methods). **(C)** Top hallmark pathways sorted by Normalized Enrichment Score per subclusters of the primary and metastatic HNSCC tumor cells (Methods). **(D)** Spatial activity of reactome programs in the primary tissue, depicted as AUCell z-scores higher than 2 (Methods) ^44^. **(E)** As in **(D)**, but for the metastatic lymph node tissue. **(F)** Gene expression enrichment in cells located at the tumor-immune boundary of metastatic lymph node section 4. Left: the defined boundary at the lymph node side (Methods); center: aggregated expression score of the top-50 genes, sorted by boundary enrichment values (*I*\*); right: sorted rank of boundary enrichment values, across the 9,000 genes with highest mean expression. **(G)** As in **(F)**, but for cells residing in the tumor side of the tumor-immune boundary.

**Figure S7:**
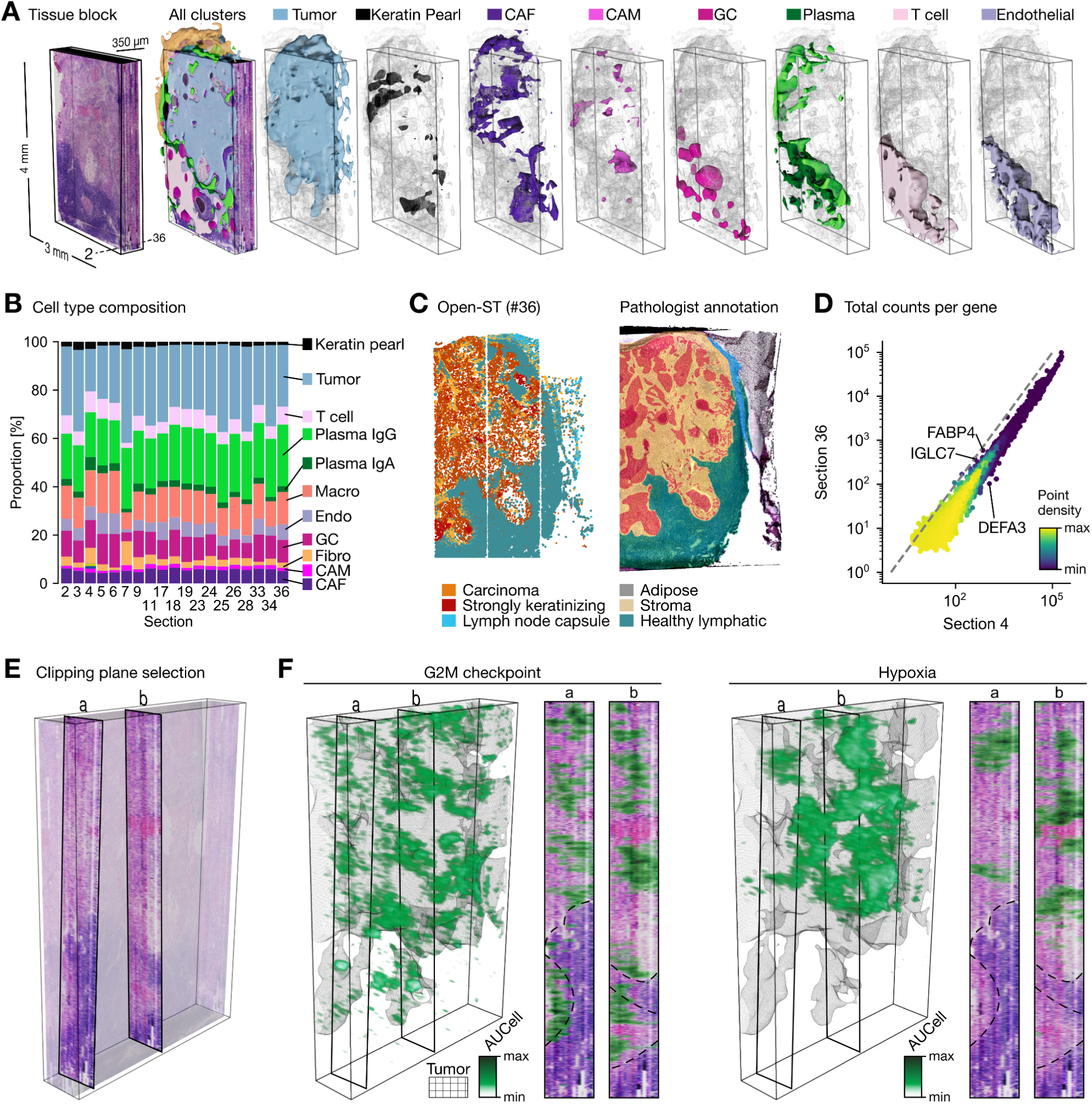
Dissecting the multi-modal 3D virtual tissue block, related to Figure 6. **(A)** 3D-rendering of tumor, stromal, and immune clusters as smooth surfaces overlaid on the virtual tissue block. Remaining clusters are shown as skeletal representation (wireframe) in the background. **(B)** Cell type composition across the 19 sections of the metastatic lymph node. **(C)** Tissue domains of a section as identified by Open-ST (left) and manually annotated by the pathologist, superimposed on H&E image (right). Open-ST clusters were merged for comparing to pathologist-annotated domains: carcinoma (tumor), strongly keratinizing (keratin pearl), stroma (CAF, CAM), healthy lymphatic (T-cells, IgA/IgG plasma cells, macrophages, endothelial, germinal center, Mast cells, cytotoxic T-cell, CCL21-expressing), lymph node capsule (fibroblasts), adipose (adipocytes). **(D)** Correlation of gene expression between section 4 and section 35 for cell type markers illustrating the relationship of mean gene expression levels (depth-corrected and log-normalized) across cells of spatially distant sections. Diagonal dashed line: y=x. **(E)** Two clipping planes orthogonal to the cutting direction (*a*, *b*) taken from the staining channel, used to project gene set activity and gene expression. **(F)** Spatial activity of the G2M checkpoint and hypoxia hallmark gene sets, quantified per segmented cell with AUCell and visualized as smoothed volumetric renderings ^44^. Tumor surface is shown as a skeletal representation (wireframe) in gray. Macro: macrophage; IFNr: interferon production regulator; Endo: endothelial; CAM: cancer-associated macrophage; CAF: cancer-associated fibroblast; GC: germinal center; Fibro: fibroblast; Cyto: cytotoxic; M1: M1 macrophage; Fibro: fibroblast; Endo: endothelial

