## Supplemental Information for "Open-ST: High-resolution spatial transcriptomics in 3D"

**Supplemental information 1**
NovaSeq6000_S4_barcoding_seq_recipe.xml

**Supplemental tables**

- **Table S1:** RNA quality and mapping statistics per sample
- **Table S2:** Global and per-cell summaries, per sample
- **Table S3:** List of marker genes for the E13 mouse head
- **Table S4:** List of marker genes for the HNSCC sample
- **Table S5:** List of marker genes for the metastatic lymph node
- **Table S6:** List of marker genes for the healthy lymph node
- **Table S7:** Bulk differential expression between tumor cells in the primary HNSCC vs. metastatic lymph node.

**Supplemental notes**

- **Supplemental note 1:** The mutual contamination of markers from pairs of cell types is local and proportional to their spatial proximity.
- **Supplemental note 2:** Spacemake yaml config file of the benchmark

**Supplemental note 1:** the mutual contamination of markers from pairs of cell types is local and proportional to the spatial proximity.

To evaluate the impact of spatial biases of Open-ST data on the transcriptomic profiles of identified cell types, especially in spatial locations with high transcriptomic diversity, we quantified crosstalk in the E13 mouse head.

We assume all (segmented) cells as the sources of transcriptomic information in the dataset, whose location in the transcriptomic manifold can be clustered into distinct populations or types. Thus, we define crosstalk as the abundance of marker genes from population $a$ in population $b$, which may arise from lateral diffusion, contributions from out-of-plane cells, or extracellular RNA mislocalized inside cell segmentation masks. Therefore, in datasets with perfect segmentation and no prior local biases in transcript capture, we expect to find no signal of the program A at cell $b$, and vice versa; that is, a scenario comparable to scRNA-seq data.

Specifically, we focused on the 'Blood' and 'Chondrocyte' clusters, as their marker genes were mutually exclusive, and pairs of cells could be found at a variety of distances. Within a proximity of 20 µm or less, we observed a higher degree of mixing of cell signatures than at greater distances, suggesting the existence of local biases in transcript capture, which may lead to unexpected gene expression patterns. Specifically, cells labeled as 'Chondrocyte', characterized by markers *Col2a1*, *Col9a1*, *Col11a1*, *Sox9* and *Col9a3*, often exhibited expression of markers *Hbb-y*, *Hbb-bt*, *Hbb-bs*, *Hba-a1* and *Hba-a2* (associated with cluster 'Blood') when in close spatial proximity to cells annotated as 'Blood' (Supplementary Figure S3D). We validated our quantification of crosstalk in two ways. Simulated diffusion of transcripts resulted in the merging of distinct cell populations, even at larger distances. Moreover, using a regular hexagonal grid as a segmentation mask led to the emergence of a mixed state corresponding to the merging of two cells under the same spatial location (Figure S3E).

These findings emphasize that our method may encounter limitations when dealing with closely spaced cell populations, where segmentation is more challenging and lateral diffusion can have a stronger effect. Spatial bias can impact current dimensionality reduction and normalization approaches, as borrowed from single-cell analyses toolboxes [1–3]. Assuming that the cell type labels are accurate, the UMAP visualization of *Hbb-y* expression reveals a gradient within the isolated 'Chondrocyte' cluster that is inversely proportional to the spatial distance to the nearest cell of the 'Blood' cluster (Supplementary Figure S3F). According to common single-cell practices, this could be interpreted as a trajectory between two adjacent and connected clusters, which would be meaningless in this scenario.

We hypothesize that the inferred normalization and expression manifold is biased by the local gradients. Therefore, caution must be taken when performing analysis relying on manifold structure, such as cell typing, pseudotime and RNA velocity, or analyses relying on counts, such as single-cell or pseudo-bulk differential gene expression.

### **Supplemental note 2:** Spacemake yaml config file of the benchmark


```
adapters:
  optical_primer: GAATCACGATACGTACACCA
  smart: AATGATACGGCGACCACCGAGATCTACACTCTTTCCCTACACGACGCTCTTC
barcode_flavors:
  default:
        UMI: r1[12:20]
        bam_tags: CR:{cell},CB:{cell},MI:{UMI},RG:{assigned}
        cell: r1[0:12]
  sc_10x_v2:
        UMI: r1[16:26]
        bam_tags: CR:{cell},CB:{cell},MI:{UMI},RG:{assigned}
        cell: r1[0:16]
  seq_scope:
        UMI: r2[0:9]
        bam_tags: CR:{cell},CB:{cell},MI:{UMI},RG:{assigned}
        cell: r1[0:20]
  slide_seq_14bc:
        UMI: r1[14:23]
        bam_tags: CR:{cell},CB:{cell},MI:{UMI},RG:{assigned}
        cell: r1[0:14]
  slide_seq_15bc:
        UMI: r1[15:23]
        bam_tags: CR:{cell},CB:{cell},MI:{UMI},RG:{assigned}
        cell: r1[0:14]
  visium:
        UMI: r1[16:28]
        bam_tags: CR:{cell},CB:{cell},MI:{UMI},RG:{assigned}
        cell: r1[0:16]
  openst:
        UMI: r2[0:9]
        bam_tags: CR:{cell},CB:{cell},MI:{UMI},RG:{assigned}
        cell: r1[2:27]
  stereo_seq:
        UMI: r1[25:35]
        bam_tags: CR:{cell},CB:{cell},MI:{UMI},RG:{assigned}
        cell: r1[0:25]
external_bin:
  dropseq_tools: ../Drop-seq_tools-2.5.1
microscopy_out: ''
puck_data:
  barcode_file: predictions_ml.csv
  root: puck_data
pucks:
  default:
        coordinate_system: ''
        spot_diameter_um: 10
        width_um: 3000
  novaseq_S4:
        coordinate_system: puck_data/novaseq_S4_coordinate_system.csv
        spot_diameter_um: 0.6
        width_um: 1200
  novaseq_S4_fc1:
```

```
        coordinate_system: puck_data/novaseq_S4_coordinate_system_fc_1.csv
        spot_diameter_um: 0.6
        width_um: 1200
  novaseq_S4_fc2:
        coordinate_system: puck_data/novaseq_S4_coordinate_system_fc_2.csv
        spot_diameter_um: 0.6
        width_um: 1200
  seq_scope:
        coordinate_system: 'puck_data/seq_scope_coordinate_system.csv'
        spot_diameter_um: 1
        width_um: 1000
  seq_scope_no_cs:
        coordinate_system: ''
        spot_diameter_um: 1
        width_um: 1000
  slide_seq:
        coordinate_system: ''
        spot_diameter_um: 10
        width_um: 3000
  slide_seq_benchmark:
        coordinate_system: ''
        spot_diameter_um: 10
        width_um: 2000
  visium:
        #barcodes: puck_data/visium_barcode_positions.csv
        coordinate_system: ''
        spot_diameter_um: 55
        width_um: 6500
  stereo_seq:
        coordinate_system: 'puck_data/stereo_seq_coordinate_system.csv'
        spot_diameter_um: 0.5
        width_um: 7000
  stereo_seq_no_cs:
        coordinate_system: ''
        spot_diameter_um: 0.5
        width_um: 7000
root_dir: ''
run_modes:
  default:
        clean_dge: false
        count_intronic_reads: true
        count_mm_reads: false
        detect_tissue: false
        mesh_data: false
        mesh_spot_diameter_um: 55
        mesh_spot_distance_um: 100
        mesh_type: circle
        n_beads: 100000
        polyA_adapter_trimming: true
        spatial_barcode_min_matches: 0
        umi_cutoff:
        - 100
        - 300
        - 500
  scRNA_seq:
```

```
        count_intronic_reads: true
        count_mm_reads: false
        detect_tissue: false
        n_beads: 10000
        umi_cutoff:
        - 500
    seq_scope:
        clean_dge: false
        count_intronic_reads: true
        count_mm_reads: false
        detect_tissue: false
        mesh_data: true
        mesh_spot_diameter_um: 10
        mesh_spot_distance_um: 15
        mesh_type: hexagon
        n_beads: 1000
        umi_cutoff:
        - 100
        - 300
    slide_seq:
        clean_dge: false
        detect_tissue: false
        n_beads: 100000
        umi_cutoff:
        - 50
    visium:
        clean_dge: false
        count_intronic_reads: true
        count_mm_reads: false
        detect_tissue: true
        n_beads: 10000
        umi_cutoff:
        - 1000
    openst:
        clean_dge: false
        count_intronic_reads: true
        count_mm_reads: false
        detect_tissue: false
        mesh_data: true
        mesh_spot_diameter_um: 10
        mesh_spot_distance_um: 10
        mesh_type: hexagon
        n_beads: 2500000
        polyA_adapter_trimming: true
        spatial_barcode_min_matches: 0.3
        umi_cutoff:
        - 5
        - 20
        - 50
  openst_mesh_7um:
        mesh_spot_diameter_um: 7
        mesh_spot_distance_um: 7
        n_beads: 100000
        spatial_barcode_min_matches: 0.3
        parent_run_mode: openst
```

benchmark_openst:
spatial_barcode_min_matches: 0.1
parent_run_mode: openst_mesh_7um
benchmark_seqscope:
spatial_barcode_min_matches: 0.1
parent_run_mode: openst_mesh_7um
benchmark_visium:
mesh_data: false
n_beads: 10000
parent_run_mode: openst
benchmark_slideseq:
mesh_data: false
n_beads: 100000
parent_run_mode: openst
benchmark_stereoseq:
spatial_barcode_min_matches: 0.05
parent_run_mode: openst_mesh_7um
species:
human:
genome:
annotation:
/data/rajewsky/home/dleonpe/projects/openst_paper/data/0_genomes/gencode.v43.basic.annotation.gt
f
sequence: /data/rajewsky/genomes/GRCh38.p13/GRCh38.p13.fa
phiX:
annotation: ''
sequence: /data/rajewsky/genomes/phiX/phiX.fa
rRNA:
annotation: ''
sequence: /data/rajewsky/projects/fc/species_data/human/rRNA/sequence.fa
mouse:
genome:
annotation: /data/rajewsky/projects/fc/species_data/mouse/genome/annotation.gtf
sequence: /data/rajewsky/projects/fc/species_data/mouse/genome/sequence.fa
phiX:
annotation: ''
sequence: /data/rajewsky/genomes/phiX/phiX.fa
rRNA:
annotation: ''
sequence: /data/rajewsky/projects/fc/species_data/mouse/rRNA/sequence.fa
temp_dir: /tmp
